# ANKRD26 is a new regulator of type I cytokine receptor signaling in normal and pathological hematopoiesis

**DOI:** 10.1101/2022.09.01.506160

**Authors:** Francesca Basso-Valentina, Alessandro Donada, Vladimir T Manchev, Manuel Lisetto, Nathalie Balayn, Jean Edouard Martin, Delphine Muller, Cecilia Paola Marin Oyarzun, Hélène Duparc, Brahim Arkoun, Alessandro Cumin, Lionel Faivre, Nathalie Droin, Ida Biunno, Alessandra Balduini, Najet Debili, Iléana Antony-Debré, Caroline Marty, William Vainchenker, Isabelle Plo, Remi Favier, Hana Raslova

**Author notes:** **Corresponding author:** Hana Raslova, INSERM UMR1287, Gustave Roussy, 114 rue Edouard Vaillant, 94805, Villejuif cedex, France. these authors contributed equally to this work (Francesca Basso-Valentina, Alessandro Donada).

## Abstract

Sustained ANKRD26 expression associated with germline *ANKRD26* mutations causes Thrombocytopenia 2 (THC2), an inherited platelet disorder associated with leukemia predisposition. Some of those patients present also erythrocytosis and/or leukocytosis. Using multiple human-relevant *in vitro* models (cell lines, primary patient cells and patient-derived iPSCs) we demonstrate for the first time that ANKRD26 is expressed during the early steps of erythroid, megakaryocyte and granulocyte differentiation, and is necessary for progenitor proliferation. As differentiation progresses, ANKRD26 expression is progressively silenced, to complete the cellular maturation of the three myeloid lineages. In primary cells, abnormal ANKRD26 expression in committed progenitors directly impacts the proliferation/differentiation balance for these three cell types. We show that ANKRD26 interacts with and crucially modulates the activity of MPL, EPOR and G-CSFR, three homodimeric type I cytokine receptors that regulate blood cell production. Higher than normal levels of ANKRD26 prevent the receptor internalization, which leads to increased signaling and cytokine hypersensitivity. Altogether these findings show that ANKRD26 overexpression or the absence of its silencing during differentiation are responsible for myeloid blood cell abnormalities in THC2 patients.

## INTRODUCTION

*ANKRD26* is the ancestor of a family of primate-specific genes termed POTE (Prostate-, Ovary, Testis-, and placenta-Expressed genes), whose gene expression is restricted to a few normal tissues and a larger number of pathological tissues. It encodes for a protein of 192 kDa, containing both spectrin helices and ankyrin repeats, protein domains known to interact with cytoskeletal and signaling proteins^1-3^. Germline mutations in the regulatory region of the gene *ANKRD26* are associated with thrombocytopenia 2 (THC2).^4^ All THC2 patients display moderate thrombocytopenia and mild bleeding, while a smaller number of patients present erythrocytosis and/or leukocytosis.^5^ Importantly, 10% of THC2 patients develop myeloid malignancies, classifying THC2 as an inherited thrombocytopenia predisposing to leukemia^5-7^. We have previously demonstrated that ANKRD26 is indeed involved in modulating TPO-dependent signaling^8^ and its expression increases the intensity of the MAPK/ERK1/2 activation. Coupled with clinical observations, we made the hypothesis that ANKRD26 plays a broader role in hematopoiesis by modifying the cytokine-mediated cell signaling.

Type I receptors, particularly the homodimeric ones, play a fundamental role in blood myeloid cell production. G-CSFR activation drives neutrophil differentiation^9^, EPO/EPOR signaling is indispensable for red blood cell production^10^, and TPO/MPL signaling is essential for megakaryocyte (MK) differentiation^11^ and hematopoietic stem cell quiescence^12^. Loss- or gain-of-function mutations of the homodimeric receptors are associated with several inherited or acquired diseases^13-16^, as are mutations in proteins directly modulating ligand/receptor signaling^17^.

These receptors are traditionally activated through ligand binding, and their cell surface density is tightly regulated to prevent aberrant activation^18^. After ligand binding, JAKs activate and phosphorylate the receptor cytoplasmic domain. This phosphorylation allows the recruitment of several signaling molecules such as the Signal Transducer and Activator of Transcription (STAT) proteins, which in turn are phosphorylated by JAKs^19,20^. Other pathways commonly activated by type I cytokine receptors, which are also dependent on JAK activation, are the Phosphoinositide 3-kinase (PI3K)/AKT pathway and the Extracellular signal-Regulated Kinase (ERK) pathway, both of which are essential for the control of cell proliferation, cell survival and differentiation^21,22^. To protect cells from over-stimulation, several mechanisms exist to attenuate signal transduction: recruitment of tyrosine phosphatases to inactivate JAKs^13^; synthesis of Suppressor of Cytokine Signaling (SOCS) proteins that can inactivate JAKs^23^; and finally receptor internalization and subsequent degradation through the proteasome and lysosome pathways^24,25^. There is therefore a strong interest in identifying the proteins able to modulate the intensity of these crucial signaling axes. Using different human-relevant models, we identified a crucial role for ANKRD26 in modulating TPO-, EPO- and G-CSF-dependent cell signaling, and thereby in the generation of the hematopoietic cells controlled by these cytokines.

## MATERIAL AND METHODS

### Study approval

Blood samples from patients and healthy subjects were collected after informed written consent and obtained in accordance with the Declaration of Helsinki. The study was approved by the Comité de Protection des Personnes CPP N°2020T2-02 and by AP-HP, Hôpital Saint-Louis, Unité de Thérapie Cellulaire, CRB-Banque de Sang de Cordon, Paris, France, N°AC-2016-2759.

Animal experiments were performed in accordance with 2010/63/UE European legislation and decree n°2013-118 of French legislation and recorded under protocol number APAFIS# 2016-008-7175.

### Primary cell culture

CD34^+^ cells were isolated from umbilical cord blood or peripheral blood by positive selection using immunomagnetic bead cell-sorting system (AutoMacs; Miltenyi Biotec).

### UT7 cell line

Human UT7 megakaryoblastic cells expressing respectively HA_MPL^26^ and HA_EPOR^27^ were previously reported, and human UT7 line expressing HA_G-CSFR was obtained by the transduction of UT7 parental cells with a retrovirus encoding HA_G-CSFR.

### iPSCs generation and expansion

Patient CD34^+^ cells were expanded in serum-free medium containing EPO (1 U/mL), FLT3l (10 ng/mL), G-CSF (20 ng/mL), IL-3 (10 ng/mL), IL-6 (10 ng/mL), SCF (25 ng/mL), TPO (10 ng/mL) and GM-CSF (10 ng/mL) for 6 days and transduced with VSV-G pseudotyped retroviruses encoding for the OSKM combination (OCT4, SOX2, KLF4 and c-MYC). Colonies with an ES-like morphology were manually isolated, expanded for a reduced number of passages and frozen. The iPSC control cell lines C1, C2 and C3 were already established and previously characterized. ^28-30^

### Clonogenic potential of primary cells in semi-solid culture

#### Methylcellulose culture assay

hematopoietic progenitors (CD34^+^CD43^+^ for iPSC and CD34^+^ for primary cells) were plated in human methylcellulose medium H4434 (STEMCELL technologies), containing recombinant human cytokines and scored for the presence of colonies 14 days after.

#### Fibrin clot assay

To assess the CFU-MK potential, CD34^+^GFP^+^ or CD34^+^Cherry^+^ sorted cells were seeded at 1.5 × 10^3^ cells/mL in triplicate in fibrin clot medium, as previously described.^29^

Other protocols are detailed in Supplementary Material and Methods and all the primers, antibodies and other reagents are listed in **Supplementary Table 1**.

### Statistics

All data are shown as mean±SD unless differentially specified in the legend. The statistical analyses were performed using the PRISM software. Statistical significance was established using a Student’s *t* test specified in legends or a One-Way Analysis of Variance (ANOVA), followed by All Pairwise Multiple Comparison Procedures. Differences were considered significant at *P* < 0.05.

## RESULTS

### ANKRD26 regulates the early steps of megakaryopoiesis

We have previously shown that the overexpression of ANKRD26 in THC2 patients megakaryocytes (MK) leads to thrombocytopenia, due to the deregulation of the TPO/MPL signaling^8^ during the late stage of differentiation. Indeed, the abnormal presence of ANKRD26 in the terminal phases of MK maturation, and the consequent stronger MAPK-pathway activation, cause a strong reduction in the number of proplatelet-forming MKs in those patients. To assess the impact of this protein on the early steps of megakaryopoiesis, we have transduced cord blood (CB)-CD34^+^ progenitor cells with a lentiviral vector over-expressing ANKRD26 and cultured them *ex vivo*. ANKRD26 overexpression only slightly increased the number of MK progenitors in presence of 10 ng of TPO, without affecting their proliferation (**Figure 1A, B, Supplementary Figure 1A**). We confirmed the previously observed reduced proplatelet formation (**Figure 1C**), without altering the MPL expression level at the MK cell surface (**Figure 1D**).

**Figure 1:**
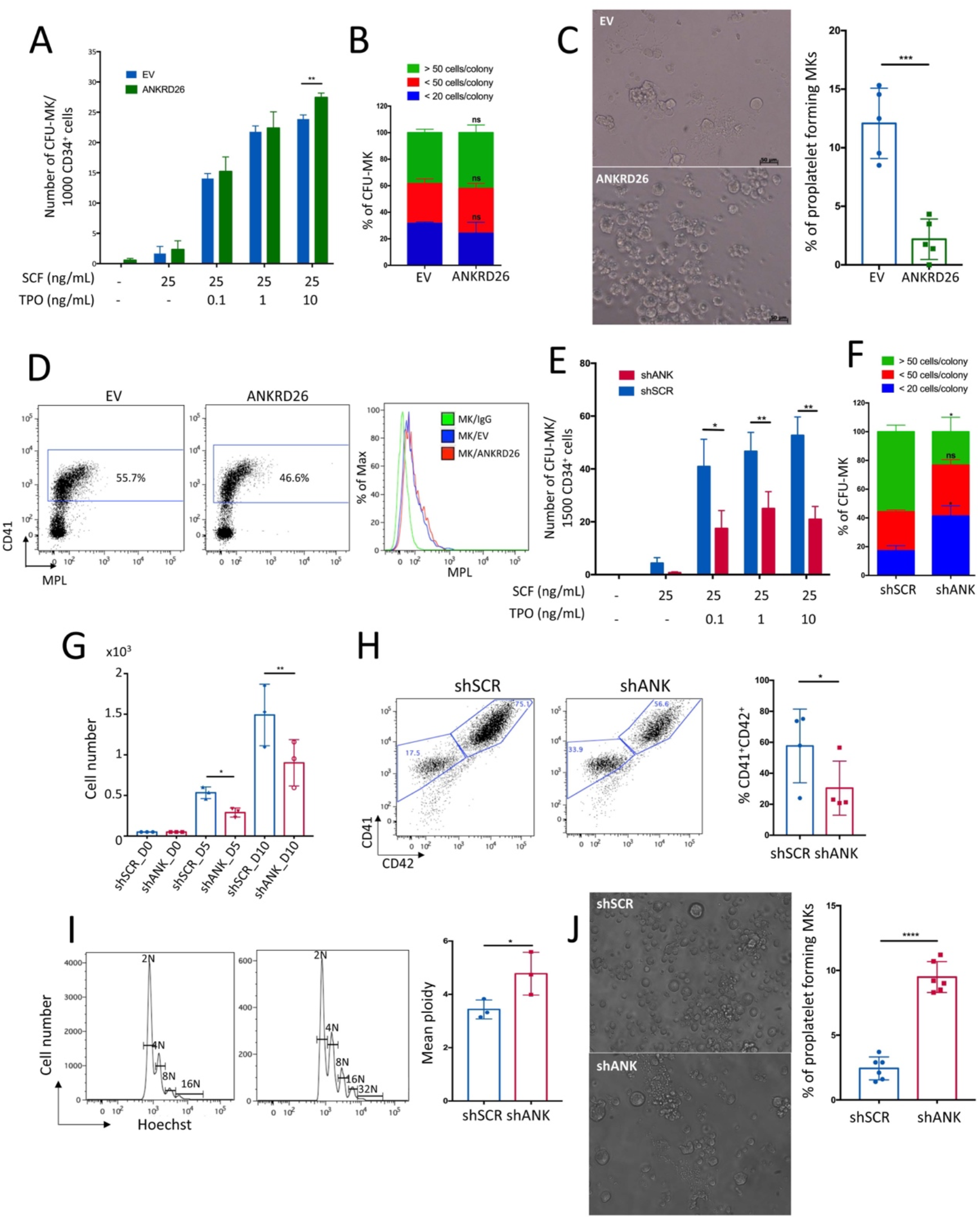
ANKRD26 regulates the early and late stages of megakaryopoiesis. A-D) Primary CD34^+^ cells were transduced with empty lentivirus (EV) or lentivirus encoding *ANKRD26_V5* and Cherry. A-B) CD34^+^-Cherry cells were sorted at day 2 after transduction and cultured in semi-solid medium (fibrin clot medium) in the presence of SCF and TPO. Colonies derived from MK-progenitors were assessed by anti-CD41 antibody labeling. B) Plating efficiency of MK-progenitors (CFU-MK) was only slightly increased after ANKRD26 overexpression in the presence of 10 ng/mL of TPO. B) the proliferative rate of CFU-MK was not affected by the increased ANKRD26 level. Shown are averages of 3 independent experiments as mean±SEM. **P<0.01; paired t-test, ns: non-significant. C-D) Primary CD34^+^ cells were transduced with empty lentivirus (EV) or lentivirus encoding *ANKRD26_V5* and gene resistant to hygromycin B. C) ANKRD26 overexpression in primary CD34^+^ cells cultured in the presence of SCF and TPO and hygromycin B prevented proplatelet formation, evaluated by inverted light microscope at day 14 of culture. Shown are averages of 5 independent experiments as mean±SD. ***P<0.005; paired t-test. D) MPL level at megakaryocytes (CD41^+^ cells) cell surface was not affected by ANKRD26 overexpression. E-J) CD34^+^ cells were transduced with shSCR or shANK and GFP encoding lentiviruses, sorted 2 days after transduction and cultured in semi-solid medium (fibrin clot medium) in the presence of SCF and TPO (E-F) or in liquid medium (G-J). E-F) Colonies derived from MK-progenitors were assessed by anti-CD41 antibody. E) Plating efficiency of megakaryocyte progenitors (CFU-MK) was decreased after inhibition of ANKRD26 (shANK), in the presence of different TPO doses. F) The proliferation rate of MK-progenitors was decreased after ANKRD26 inhibition, as shown by the increase of colonies composed of less than 20 cells and the decrease of those with more than 50 cells. Shown are averages of 3 independent experiments as mean±SEM. *P<0.05; **P<0.01; ns: non-significant; paired t-test. G-H) The inhibition of ANKRD26 led to a decrease in the proliferation rate in liquid culture (G) and to a decreased frequency of MKs (CD41^+^CD42^+^ cells) at day 10 of culture (H). I-J). In contrast, an increase in ploidy level (I) and in the percentage of proplatelet-forming MKs was detected after ANKRD26 inhibition (J). Shown are averages of 3 (G, I), 4 (H) or 6 (J) independent experiments as mean±SD, *P<0.05; ****P<0.001, paired t-test. All the experiments of ANKRD26 inhibition were performed at least twice with shANK_1 and once with shANK_2.

ANKRD26 has been shown to be more expressed in CD34^+^ progenitor cells than in mature MKs, suggesting that its expression is necessary during the early stages of hematopoiesis^8,31^ (**Supplementary Figure 2A, B)**. As overexpression had no major effect at this stage, we decided to investigate if ANKRD26 down-regulation at the CD34^+^ progenitor level could affect the number of MK progenitors (CFU-MK). We transduced CB-CD34^+^ hematopoietic progenitors with lentiviruses encoding short hairpin RNAs (sh) against *ANKRD26* (shANK_1 or shANK_2) (**Supplementary Figure 1B**) and cultured them in fibrin clot medium, in presence of stem cell factor (SCF) and increasing doses of TPO. We observed a 50% decrease in CFU-MK plating efficiency (**Figure 1E**), as well as a reduction of their proliferative rate, as demonstrated by the lower number of MKs per colony (**Figure 1F**). This result was confirmed in liquid culture in presence of SCF and TPO (**Figure 1G, Supplementary Figure 1C**). Interestingly, the ANKRD26 reduction led also to a reduced frequency of mature MKs (CD41^+^CD42^+^) at day 10 of culture (**Figure 1H**), but those cells displayed higher ploidy level (**Figure 1I**) and enhanced proplatelet formation capacity (**Figure 1J**).

Altogether these results demonstrate that ANKRD26 is required for optimal proliferation and differentiation of MK progenitors, but that its downregulation is required for thrombogenesis.

### ANKRD26 regulates the early steps of granulopoiesis

ANKRD26 expression decreases not only during normal MK but also during erythroid and granulocytic differentiation **(Supplementary Figure 2A, B**). To confirm that ANKRD26 deregulation could be responsible for the leukocytosis detected in some THC2 patients, we studied the granulopoiesis in primary patient cells (**Supplementary Table 2**). Although the ANKRD26 expression was not different in early progenitors (CD34^+^ cells) isolated from the peripheral blood of THC2 patients, a significant increase in ANKRD26 levels was detected during *in vitro* granulocyte differentiation (**Figure 2A**). To evaluate the biological effects of increased ANKRD26 expression, we assessed the colony forming potential of patients CD34^+^ cells in semi-solid medium and their proliferative rate in presence of SCF, IL-3 and G-CSF^32^. We detected a more than two-fold increase in granulocyte colony (CFU-G) numbers (**Figure 2B**), as well as an increased proliferative capacity (**Figure 2C**), compared to healthy controls. We performed May-Grünwald Giemsa (MGG) staining at day 15 of culture for two patients, and we detected an increased frequency of immature cells (myeloblasts, promyelocytes and myelocytes) compared to controls, while the percentages of more mature metamyelocytes and polynuclear neutrophils were clearly decreased (**Figure 2D**). Nevertheless, the percentages of mature granulocytes for patient cells increased over time (**Supplementary Figure 3A**), suggesting a maturation delay due to the increased proliferation, and not a complete blocking of differentiation. Overall, these results show that the ANKRD26 overexpression leads to defective granulopoiesis in THC2 patients, with increased proliferation and delayed maturation. This could be the first step to a pre-leukemic state. Also, it should be noted that different AML subtypes express ANKRD26 **(Supplementary Figure 2B)**.

**Figure 2:**
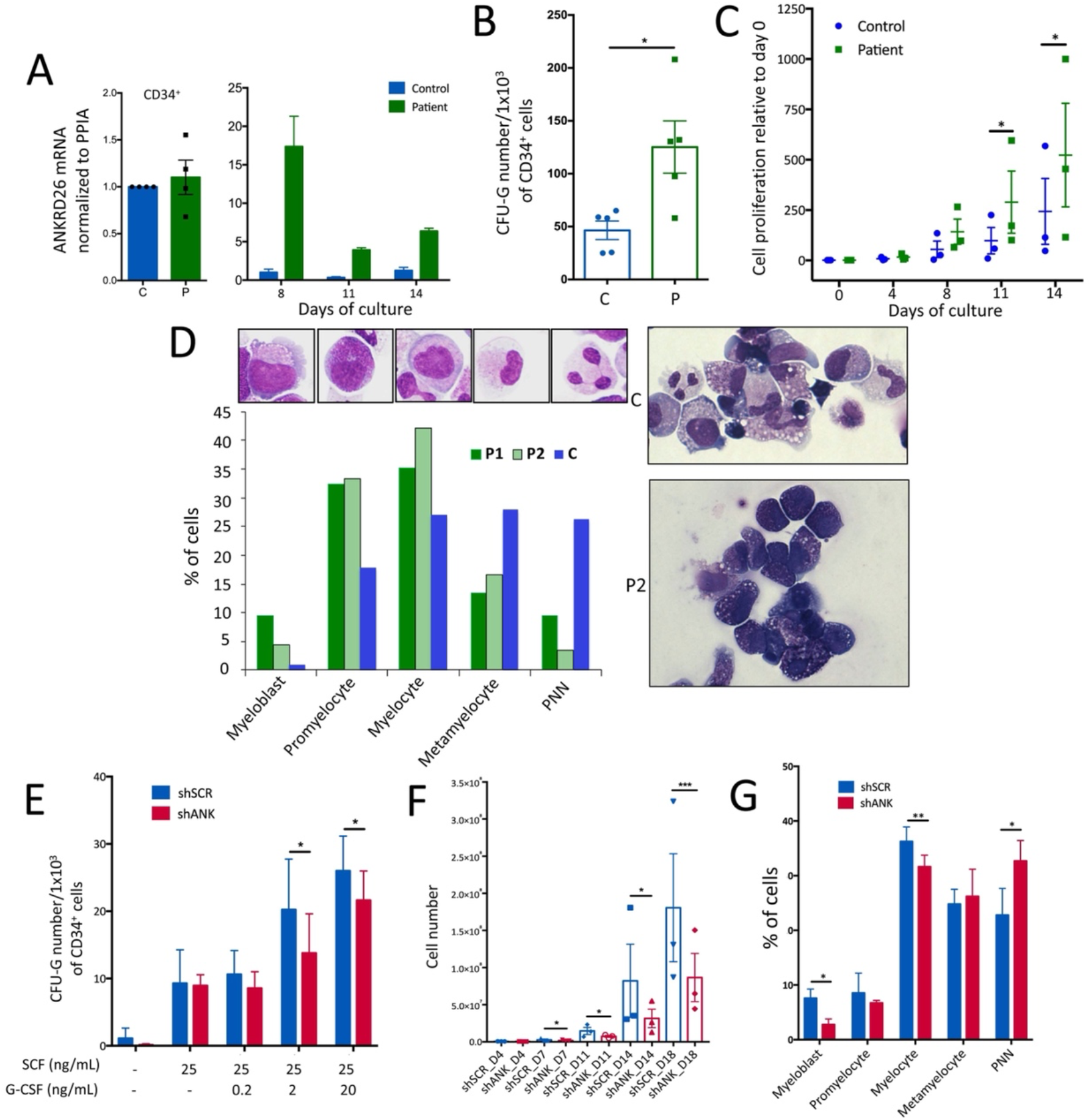
ANKRD26 regulates the early stage of granulopoiesis. Primary CD34^+^ cells obtained from peripheral blood of THC2 patients with different 5’UTR mutations were induced to granulocytic differentiation in the presence of G-CSF, IL-3 and SCF. A) ANKRD26 expression in patient CD34^+^ cells was similar to control CD34^+^ cells obtained from healthy individuals, but increased during the *in vitro* granulocytic differentiation, with a peak at day 8 of culture. *ANKRD26* transcript level was normalized to *PPIA*. Shown are averages of 4 (for CD34^+^ cells) (i) and of 2 (for granulocytic differentiation) independent experiments (ii). B) The number of patient myeloid progenitors (CFU-G) was significantly higher, compared to control progenitors, as assessed by methylcellulose assay. Shown are averages of 5 independent experiments as mean±SEM. *P<0.05; t-test with Mann Whitney correction. C) Proliferation assay performed in liquid culture supplemented with G-CSF, SCF and IL-3 showed a significant increase in cell number for patient samples at days 11 and 14 of culture. The cell count was normalized to day 0. Shown are averages of 3 independent experiments, mean±SEM, *P<0.05; *P<0.01; paired t-test. D) MGG staining of control and patient cells at different days of culture in the presence of G-CSF, SCF and IL-3. MGG staining of 2 patients and 1 control samples at day 15 of culture showed an increase in the proportion of immature cells (myeloblasts, promyelocytes and myelocytes) and a decrease in the proportion of more mature cells (metamyelocytes and polynuclear neutrophils (PNN)). E-G) Effect of ANKRD26 inhibition on granulocytic lineage. CD34^+^ cells were transduced with lentiviruses encoding respectively shSCR and shANK (shANK_1 or shANK_2. E) CD34^+^-GFP cells were sorted 2 days after transduction and grown in semi-solid medium (methylcellulose) in the presence of 25 ng/mL SCF and different doses of G-CSF. Granulocytic progenitors (CFU-G) were enumerated at day 14 of culture. F-G) CD34^+^-GFP cells were sorted 2 days after transduction and grown in liquid medium in presence of 25 ng/mL SCF, 10 ng/mL IL-3 and 20 ng/mL G-CSF. F) Proliferation assay showed a significant decrease in shANK transduced cell number at days 7, 11, 14 and 18. G) MGG staining performed at day 14 of culture showed an accelerated differentiation of shANK-transduced cells.

To further confirm the role of ANKRD26 in the granulocytic lineage, we transduced the CB-CD34^+^ progenitors with lentiviruses encoding shANK and cultured them in presence of increasing doses of G-CSF. We detected a reduction in the number of granulocytic progenitor-derived colonies (CFU-G) in methylcellulose assay in presence of 2 and 20 ng of G-CSF (**Figure 2E**). We also observed a decreased proliferation rate in liquid culture (**Figure 2F**), with no effect on apoptosis (**SF 3B**), and an accelerated differentiation (**Figure 2G**), confirming the essential role of ANKRD26 in the early stages of granulopoiesis.

### Patient-derived iPS cells recapitulate the defective granulopoiesis

To overcome the limitations associated with the rarity of THC2 patients, we derived iPSC lines from two patients harboring two different mutations: c.-118A>C (ANK1) and c.-127delAT (ANK2) (**Supplementary Table 3**). The iPSC lines were generated from CD34^+^ cells isolated from peripheral blood via integrative reprogramming and were characterized for their phenotypic and functional pluripotency (**Supplementary Figure 4**). Three different clones were studied for each patient-derived iPSC. As a control, we used three different iPSC lines derived from healthy individuals^28-30^.

We used a serum-free, xeno-free differentiation protocol (**Supplementary Figure 5**). We isolated the hematopoietic progenitors after 14 days of culture (CD34^+^CD43^+^), described to have a bias towards the granulo-monocytic differentiation output^30^. The *ANKRD26* mRNA levels in patient iPSC-derived CD34^+^CD43^+^ cells were three times higher than in controls (**Figure 3A**). This difference with respect to ANKRD26 expression in CB-derived early progenitors (CD34^+^ cells) can be explained by the fact that iPSC-derived CD34^+^CD43^+^ cells are developmentally different from adult CD34^+^ and already committed to the myeloid lineage. We detected three times more CFU-G colonies in semi-solid medium compared to the healthy controls (**Figure 3B**). An increased proliferative rate was also detected in liquid culture, in the presence of SCF, IL-3 and G-CSF (**Figure 3C**). CD15^+^ granulocytic cells derived from CD34^+^CD43^+^ patient cells expressed higher levels of ANKRD26 relative to control CD15^+^ cells (**Figure 3D**). Flow cytometry and MGG analysis displayed a delay in granulocytic differentiation at days 17-23 of culture **(Supplementary Figure 6A)**: myeloid cells derived from patient iPSC showed less than 10% of mature CD11b^+^CD15^+^CD14^-^ cells, compared to about 30% for the control cell lines at day 23 (**Figure 3E**). Overall, these results clearly highlight a defective granulopoiesis, similar to what was observed for primary cells. To bring more insights into this defect, we investigated the transcriptomic profile of the patient granulocytic progenitors: iPSC-derived granulocytic progenitors (CD43^+^CD11b^+^CD14^-^) were sorted (**Figure 3Fi**) and profiled via RNA-seq. We found 24 genes significantly upregulated and 7 down-regulated (*P*<1×10^5^) in patient cells compared to controls (**Figure 3Fii, Supplementary Table 4**). The Gene Set Enrichment Analysis (GSEA) performed with two different databases (KGEA and GOBP) revealed a significant enrichment in JAK/STAT signaling pathway in patient-derived iPSC progenitors (**Figure 3Gi, ii**) with a tendency to increased granulocytic count and delayed myeloid development (**Supplementary Figure 6B**) confirming the results obtained with primary patient cells and with shANK in CB-derived progenitors.

**Figure 3:**
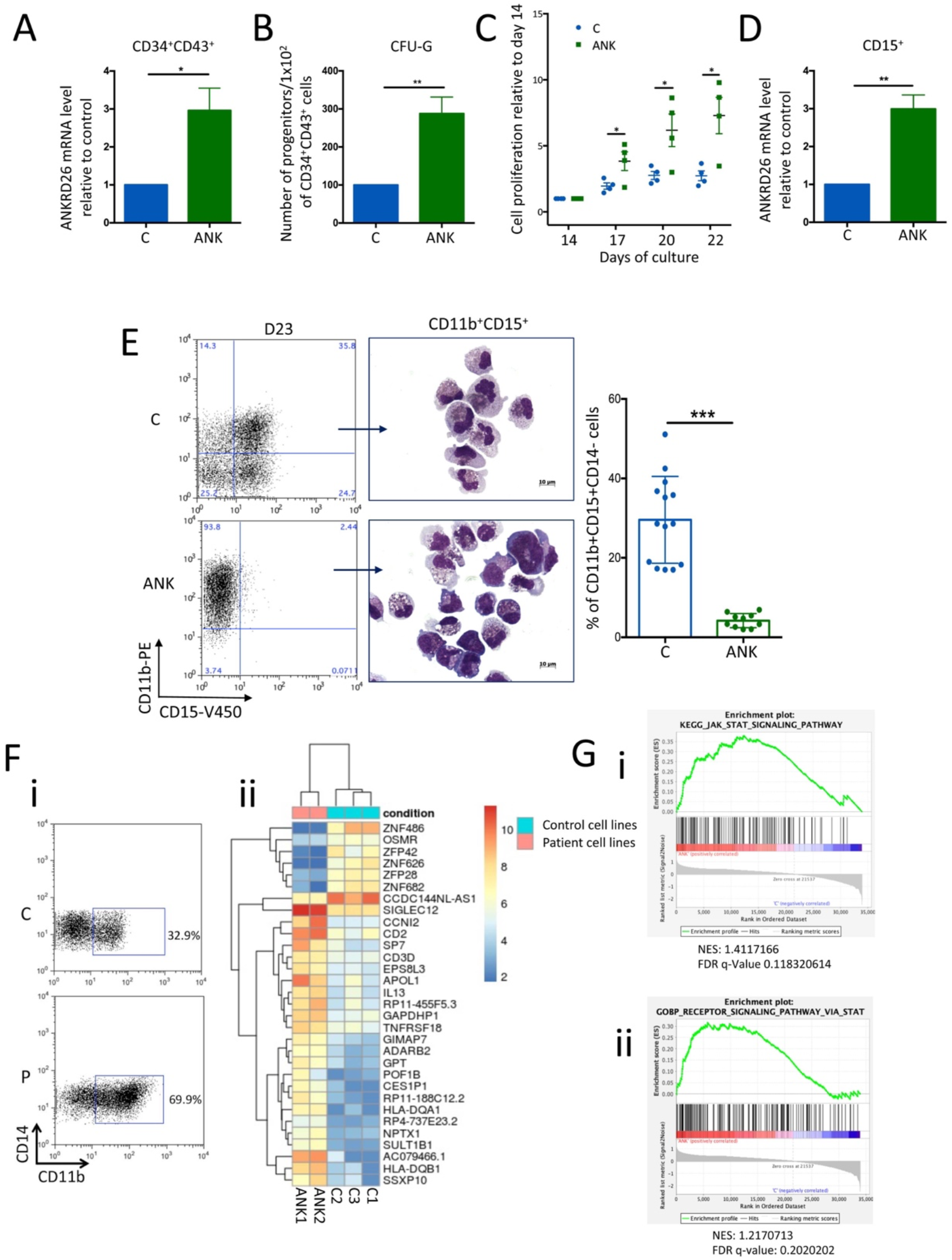
Increased ANKRD26 expression level boosts proliferation of granulocyte progenitors in an iPSC model, through enhanced JAK/STAT pathway. CD34^+^CD43^+^ progenitors derived from patient and control iPSC were differentiated as granulocytes in the presence of G-CSF, SCF and IL-3. A) ANKRD26 expression in patient CD34^+^CD43^+^ progenitors was increased when compared to controls. *ANKRD26* transcript was normalized to *PPIA*. Shown are averages of 4 independent experiments as mean±SD. *P<0.05; paired t-test. B) The number of patient granulocytic progenitors (CFU-G) derived from iPSCs was significantly higher than the control, as assessed by methylcellulose assay. Shown are averages of 7 independent experiments as mean±SD. **P<0.01; paired t-test. C) The proliferation rate was significantly increased at days 17, 20 and 22 of culture for patient cells as compared to controls. Shown are averages of 4 independent experiments as mean±SEM. *P<0.05; paired t-test. D) ANKRD26 expression in CD11b^+^CD15^+^CD14^-^ cells derived from patient iPSC was significantly increased compared to controls. *ANKRD26* transcript was normalized to *PPIA*. Shown are averages of 6 independent experiments as mean±SD. **P<0.01; paired t-test. E) CD11b^+^CD15^+^CD14^-^ cells frequency analyzed at days 21-23 reflecting a delay in maturation for patient cells as shown by FACS analysis (left panel), MGG staining (middle panel) and confirmed by statistical analyses (n=14 for control and n=10 for patient cells, mean±SEM. **P<0.005, unpaired t-test. F) WB analysis of G-CSFR/G-CSF/STAT3 signaling, performed at day 20 of culture, showed increased STAT3 phosphorylation in patient cells (ANK2) compared to control. G) RNA-seq assay was performed on CD43^+^CD11b^+^CD14^-^ progenitors sorted at day 16 of culture (Figure 5i). 7 down-regulated and 24 up-regulated genes in patient cells (n=3 for each, ANK1 and ANK2) (*P*<1×10^5^) were identified compared to controls (n=1 for C1 and C3, n=3 for C2). H) GSEA for i) KEGG_JAK_STAT_signaling pathway, for ii) for receptor signaling pathway via STAT (GO:0007259, GOBP_RECEPTOR_SIGNALING_PATHWAY_VIA_STAT) in ANK versus Control iPSC.

Within the significantly up-regulated genes, the *CCNI2* gene (Cyclin I Family member 2), a cyclin responsible for the cyclin-dependent kinase 5 (CDK5) activity, caught our attention: CDK5 was previously described to phosphorylate NOXA, promoting its cytosolic sequestration and apoptosis suppression in leukemic cells ^33^. Its expression was almost completely absent from control iPSC-derived CD43^+^CD11b^+^CD14^-^ cells. After validation of CCNI2 up-regulation in patient iPS-derived progenitors by qRT-PCR (**Supplementary Figure 6C**), we transduced the CD34^+^CD43^+^ progenitors at day 14 of culture with a lentivirus encoding shCCNI2 (shCCNI2_1 and shCCNI2_2), sorted the GFP^+^ cells 48 hours later and measured their proliferative rate at days 19, 21 and 23. The inhibition of CCNI2 by shRNA (**Supplementary Figure 6Di**) significantly reduced the number of granulocytes in patient-derived iPSCs (**Supplementary Figure 6Dii**). Although an *in silico* analysis of *CCNI2* promoter revealed a presence of 2 STAT5 binding sites (data not shown), its implication in the observed phenotype should be further investigated.

### ANKRD26 regulates the early steps of erythropoiesis

The role of ANKRD26 in the regulation of erythropoiesis was investigated in CB-CD34^+^ cells. ANKRD26 downregulation led to a reduction in the plating efficiency of erythroid progenitors (BFU-E) in semi-solid assay, in the presence of SCF and increasing doses of EPO (**Figure 4A**). Moreover, colonies from shANK progenitors had a paler color, indicating a decrease in hemoglobin content at lower EPO doses (0.01 and 0.1 U/mL), a sign of impaired differentiation (**Figure 4B**). A delayed erythroid differentiation in the presence of lower ANKRD26 levels was confirmed in liquid medium, in the presence of EPO, SCF and IL-3 (**Figure 4Ci, ii**). Finally, CB-CD34^+^ cells transduced with shANK and cultured in erythroid condition showed a reduced proliferation compared to the control (**Figure 4Di**). This is consistent with the increased proliferation observed for THC2 patient-derived cells cultured in the same condition (**Figure 4Dii**).

**Figure 4:**
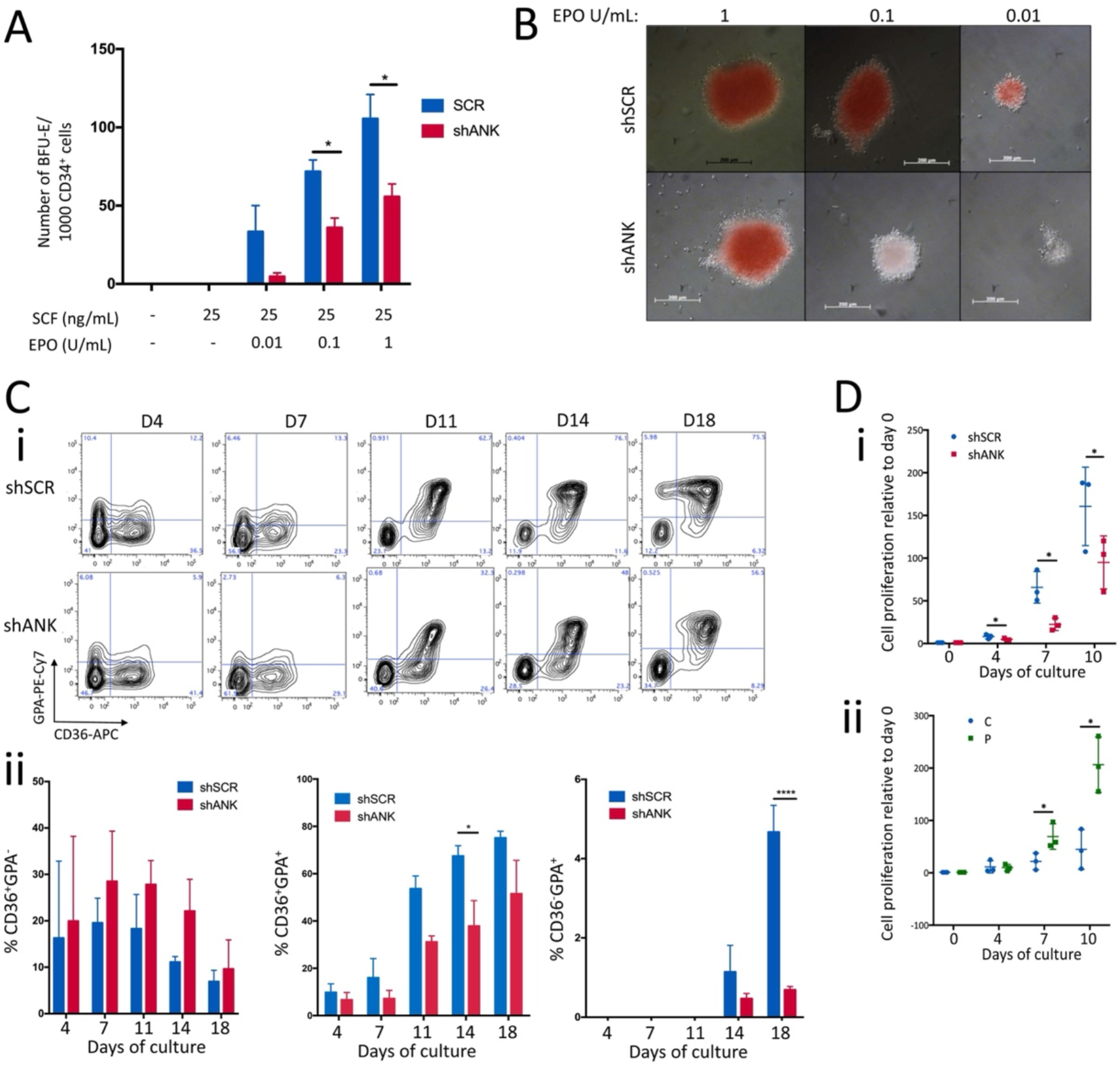
ANKRD26 regulates the early stage of erythropoiesis. A-B) CD34^+^ cells were transduced with lentiviruses encoding shSCR and shANK respectively and erythroid progenitors (BFU-E) were grown in semi-solid medium (methylcellulose) in the presence of 25 ng/mL SCF and different doses of EPO and enumerated at day 14 of culture. A) ANKRD26 inhibition led to a significant decrease in the BFU-E number. Shown are averages of 3 independent experiments as mean±SD. *P<0.05; one-tailed t-test with Mann-Whitney correction. B) Representative pictures of BFU-E colonies showing that ANKRD26 inhibition led to a lack of hemoglobinization of BFU-E derived colonies, both at 0.1 and 0.01 U/mL of EPO. C-D) Transduced CD34^+^ cells (C-Di) or primary patient CD34^+^ progenitors (Dii) were cultured in liquid medium in the presence of EPO (1 U/mL), SCF and IL-3 for 18 days. C) Kinetics of erythroid differentiation assessed by FACS (i) showed that ANKRD26 inhibition leads to a delay in differentiation. CD36^-^GPA^-^ cells represent immature, CD36^+^GPA^+^ intermediate and CD36^-^GPA^+^ mature erythroid cells. ii) Statistical analysis of different populations is shown as the average of 3 independent experiments (mean±SD, *P<0.05, ***P<0.005, 2-way Anova with multiple comparisons). D) ANKRD26 expression level affected the proliferation of CD34^+^ cells grown in erythroid conditions. i) Inhibition of ANKRD26 leads to a significant decrease in the proliferation of transduced CD34^+^ cells cultured in erythroid conditions. Shown is the average of 3 independent experiments as mean±SD, *P<0.05; paired t-test ii) Proliferation rate of primary THC2 patient CD34^+^ cells in erythroid condition was significantly higher compared to controls. Shown is the average of 3 independent experiments as mean±SD, *P<0.05; t-test with Mann-Whitney correction.

### ANKRD26 regulates MPL, G-CSFR and EPO-mediated signaling

To further understand how ANKRD26 could modify the proliferation and differentiation of hematopoietic progenitors under TPO, G-CSF and EPO stimulation, we investigated whether it may interfere with the signaling of their cognate receptors. For these studies we used two cell lines, the murine pro-B Ba/F3 and the human erythro-MK UT7 cell lines. Those cells lines were transduced with retroviruses encoding for *MPL, G-CSFR* or *EPOR*, therefore made dependent on the cognate cytokines, respectively TPO, G-CSF and EPO. In the Ba/F3 cell line, we overexpressed human ANKRD26, while in the UT7 cell line we down-regulated ANKRD26 expression by two different shRNAs, as those cells express high levels of the gene.

ANKRD26 levels did not affect MPL, G-CSFR and EPOR expression at the cell surface of UT7 at the steady state **(Figure 5A, B, C)**. We confirmed our previous observation^8^ that higher ANKRD26 levels induced stronger STAT5, ERK, AKT and also STAT3 phosphorylation in presence of 10 ng/mL of TPO (**Figure 5D, Supplementary Figure 7A, B)**. Moreover, we observed that a very low TPO dose (0.01 ng/mL) was still able to stimulate TPO/MPL signaling in cells expressing higher levels of ANKRD26 (**Figure 5E**). To better understand the interaction between MPL and ANKRD26, we assessed the receptor response to a different ligand than TPO. We used the TPO mimetic Eltrombopag (ELT), which binds the receptor at the level of the residue H499^34^, therefore causing the receptor to assume a different conformation than done by TPO^35^. Higher levels of ANKRD26 led to the same hypersensitivity to ELT as observed for TPO (**Supplementary Figure 7Ci**), suggesting that the main effect of ANKRD26 is to stabilize MPL surface expression regardless its conformation.

**Figure 5:**
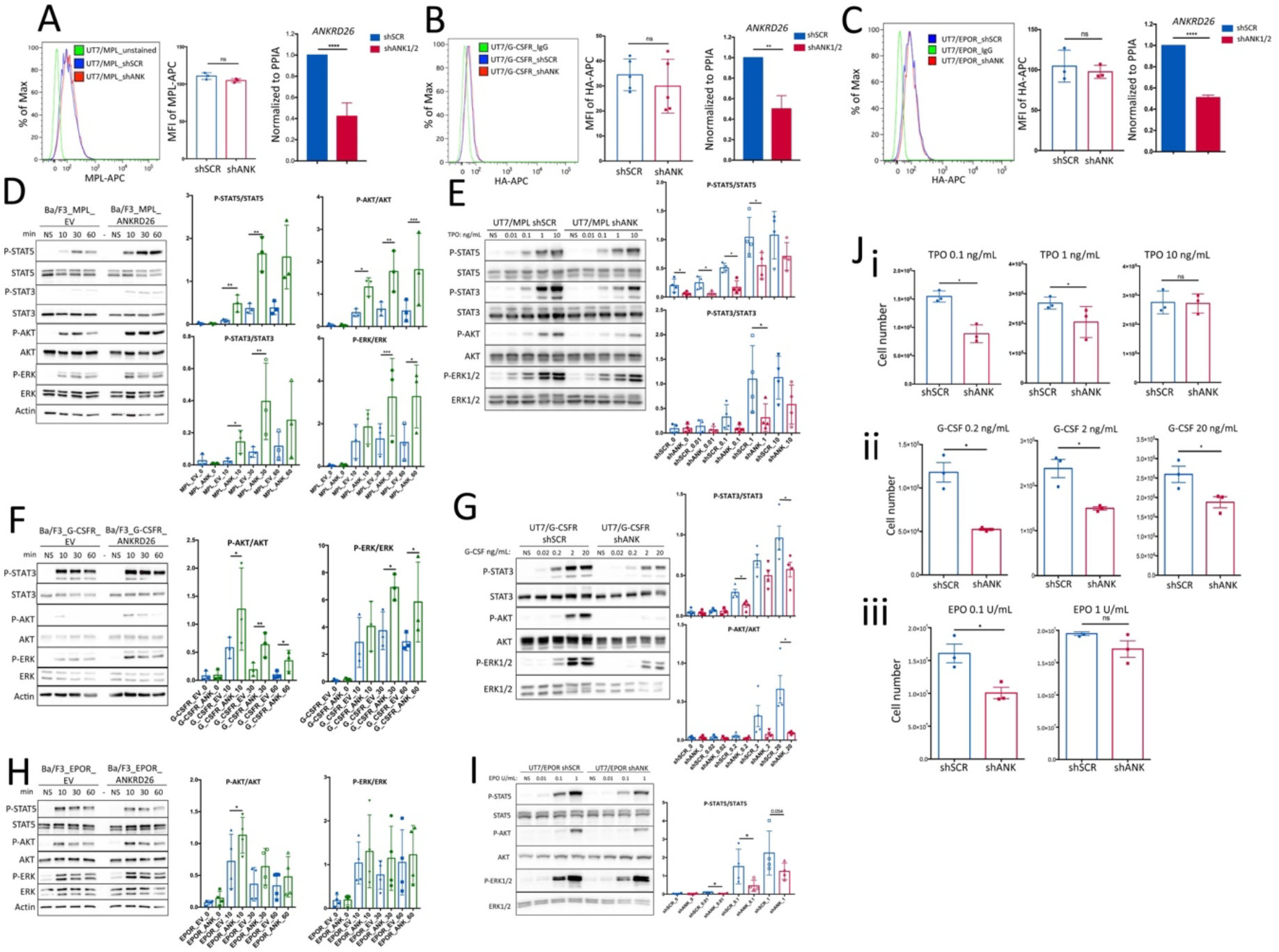
ANKRD26 regulates MPL, G-CSFR and EPO-mediated signaling. UT7 or Ba/F3 cell lines expressing respectively MPL, G-CSFR, EPOR were transduced with the lentiviruses harboring control scramble shRNA (shSCR), shANKRD26 (shANK), *ANKRD26* cDNA or empty virus (EV). A-C) Down-regulation of ANKRD26 expression level did not affect the expression of MPL measured with anti-MPL antibody (A), G-CSFR measured with anti-HA antibody (B) or EPOR measured with anti-HA antibody. The receptor levels are presented as median fluorescence intensity (MFI) at the cell surface. *ANKRD26* transcript level was normalized to *PPIA* housekeeping gene. Shown are the averages of 3 independent experiments as mean±SD (2 with shANK1_1 and 1 with shANK_2). **P<0.01; ****P<0.001; ns: non-significant, t-test with Mann Whitney correction. D-I) One of at least three independents Western Blot (WB) analyses on signaling proteins in Ba/F3 and UT7 cells, at different times post-stimulation with 10 ng/mL of TPO (D), 20 ng/mL of G-CSF (F), 1 U/mL of EPO (H), and different TPO doses (F), G-CSF doses (G), EPO doses (I) at 10 min. The histograms show quantification of the WBs representing averages of 3 or 4 independent experiments as mean±SD. *P<0.05; paired t-test. J) Number of UT7/MPL, UT7/G-CSFR, UT7/EPOR cells measured at day 4 of culture, with 3 different doses of TPO (i), 3 different doses of G-CSF (ii) and 2 different doses of EPO (iii) are shown as mean±SEM of three independent experiments, *P<0.05, ns: non-significant, t-test with Mann-Whitney correction was used.

Similarly, we showed that the higher ANKRD26 levels led to a stronger G-CSF-mediated STAT3 and AKT phosphorylation, with a similar tendency for ERK1/2 (**Supplementary Figure 8A**). The overexpression of human ANKRD26 in Ba/F3 cell line expressing G-CSFR did not affect the STAT3 activation, but led to the enhanced ERK and AKT phosphorylation (**Figure 5F, Supplementary Figure 8B**). Using UT7/G-CSFR cells, we also observed the activation of STAT3 at lower G-CSF doses (0.2 ng/mL), in the presence of higher ANKRD26 levels (**Figure 5G**).

Finally, we observed that higher ANKRD26 levels led to a stronger and more sustained EPO/EPOR-mediated activation of ERK1/2, AKT and, to a lesser extent, of STAT5 in UT7/EPOR cells (**Supplementary Figure 9A**). Increased level of AKT phosphorylation and the same tendency for ERK1/2 were also detected in Ba/F3 cell line overexpressing ANKRD26 and EPOR (**Figure 5H, Supplementary Figure 9B**). Moreover, significantly increased levels of STAT5 phosphorylation were detected at 0.1 U/mL of EPO in the presence of higher ANKRD26 levels (**Figure 5I)**.

To analyze the biological consequence of the TPO, G-SCF and EPO hypersensitivity, we measured the proliferation rate of UT7/MPL, UT7/G-CSFR and UT7/EPOR cells expressing shSCR and shANK in presence of different cytokine concentrations. We observed that for the 0.1 ng/mL TPO concentration, only the UT7/MPL cells with higher ANKRD26 levels were able to proliferate, after 4 days of culture. At higher doses of TPO (10 ng/mL), no difference in the proliferation rate was detected **(Figure 5Ji)**, a finding that corroborates the idea of receptor saturation at higher doses. A similar result was obtained in presence of ELT **(Supplementary Figure 7Cii)**. In a similar manner, UT7/G-CSFR cells expressing higher levels of ANKRD26 were able to proliferate in the presence of 0.2 ng/mL of G-CSF, contrarily to the cells expressing lower ANKRD26 levels. Using higher doses of G-CSF (2 ng/mL), ANKRD26 down-regulation led to a decreased proliferation rate, suggesting a progressive saturation of the cell surface pool of available G-CSFR (**Figure 5Jii**). Finally, UT7/EPOR cells expressing higher ANKRD26 levels proliferated more at low EPO concentration (0.1 U/mL) than the control. This proliferation gap was absent at higher EPO doses (1 U/mL) (**Figure 5Jiii)**.

Overall, these results demonstrate that higher ANKRD26 levels lead to MPL, G-SCF and EPO hypersensitivity.

### ANKRD26 interacts with the homodimeric type I receptors

We explored whether ANKRD26 was directly or indirectly affecting the receptors signaling. First, we investigated whether ANKRD26 interacted with MPL, G-CSFR and EPOR respectively. To this end, HEK293T cells were transduced with retroviruses encoding respectively HA_MPL, HA_G-CSFR or HA_EPOR and with a lentivirus encoding ANKRD26_V5. The presence of ANKRD26 in the same protein complex with each of the three receptors was demonstrated by co-immunoprecipitation assays (**Figure 6A-C**). As a proof of concept, the ANKRD26 and MPL interaction was confirmed by proximity ligation assays in UT7 cell line **(Supplementary Figure 10)** and in γ2A cells, where it was shown that JAK2 was dispensable for this interaction, although its presence slightly enhanced it (**Figure 6D**).These results demonstrate that ANKRD26 is present in the protein complex with each of these three homodimeric receptors.

**Figure 6:**
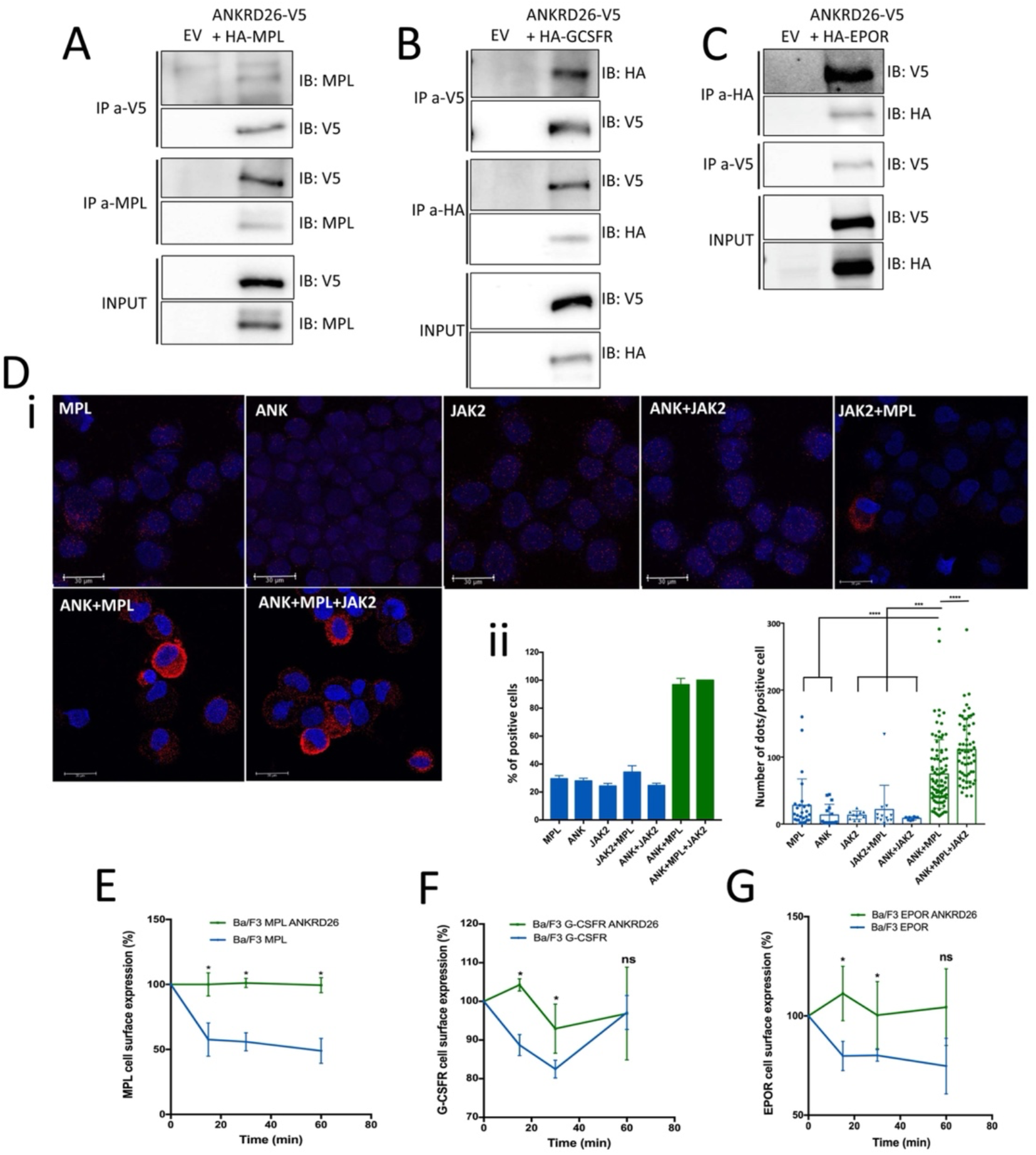
ANKRD26 interacts with homodimeric type I receptors and regulates their internalization. A-C) Co-immunoprecipitation assay performed in 293HEK cells showing the presence of ANKRD26 and MPL (A), ANKRD26 and G-CSFR (B), and ANKRD26 and EPOR (C) in the same protein complex. For each receptor, one of three independent experiments with similar results are shown. Input represents WB analysis of cells expressing empty vectors or cells co-expressing ANKRD26_V5 and HA_MPL (A), ANKRD26_V5 and HA_G-CSFR (B), ANKRD26_V5 and HA_EPOR (C). IP: immunoprecipitation, IB: immunoblot. The antibodies used were anti-V5 (for ANKRD26_V5), anti-MPL (for HA_MPL), anti-HA (for G-CSFR and EPOR). D) Proximity ligation assay for the ANKRD26 and MPL interaction. FLAG_ANKRD26 (ANK), HA_MPL (MPL) and JAK2 were overexpressed in gamma-2A cells (cells not expressing endogenous Jak2). Monoclonal anti-FLAG antibody was used for ANKRD26 and polyclonal anti-HA for MPL. i) Representative pictures of PLA for ANKRD26 and MPL interaction. The red staining represents ANKRD26/MPL interaction, scale bar = 30 ⌈m. Data in ii (left panel) represent the 2 independent experiments. At least 40 positive cells were analyzed for each condition (right panel). ***P<0.005, ****P<0.001, t-test with Mann Whitney correction. E-G) Ba/F3 cells overexpressing respectively the three receptors and transduced either with empty lentivirus or ANKRD26 cDNA encoding lentivirus were used for internalization assays. Internalization of MPL was measured with anti-MPL antibody (E), and of G-CSFR (F) and EPOR with anti-HA antibody (G). E) In the absence of ANKRD26 overexpression, almost 50% of cell surface MPL was internalized as soon as 15 min after TPO addition, while in the presence of ANKRD26, MPL was not internalized. MFI is normalized to that of Ba/F3/HA_MPL cells expressing empty vector. F) ANKRD26 overexpression inhibits G-CSFR internalization at 15 and 30 min after stimulation with 20 ng/mL of G-CSF of starved Ba/F3 cells. MFI is normalized to that of Ba/F3/HA_GCSFR cells expressing empty vector. G) ANKRD26 overexpression significantly inhibits EPOR internalization at 15 and 30 min after starved Ba/F3 cell stimulation with 1 U/mL of EPO. MFI is normalized to that of Ba/F3/HA_EPOR cells expressing empty vector. E-G) Shown are the averages of 3 independent experiments as mean±SD. *P<0.05; un-paired t-test with Mann-Whitney correction.

Furthermore, as ANK repeat-containing proteins were shown to interact with different receptors and to participate in their internalization, one possible mechanism was that ANKRD26 affects receptor-mediated signaling by changing their internalization. To investigate this hypothesis, we used the murine in Ba/F3 cell line overexpressingANKRD26 and MPL, G-CSFR or EPOR respectively, as this cell line is a better model for internalization studies than UT7 cells^36^. After overnight starvation, ANKRD26 did not modify receptor expression at cell surface **(Supplementary F 7A, 8B, 9B)**. The MPL-expressing Ba/F3 cells were then exposed to 50 ng/mL of TPO. In the absence of human ANKRD26, about 40% of MPL was internalized 15 minutes after TPO exposure and additional exposure only slightly increased the quantity of internalized MPL. In contrast, the presence of human ANKRD26 almost completely abrogated MPL internalization, even after 60 min of TPO exposure **(Figure 6E)**. Similarly, the overexpression of human ANKRD26 in BaF/3 cell line expressing G-CSFR **(Figure 6F)** or EPOR **(Figure 6G)** led to a defect in G-CSFR or EPOR internalization at 15 and 30 min of stimulation of starved cells with G-CSF and EPO respectively. These results demonstrate that receptor internalization is finely regulated by ANKRD26 levels.

## DISCUSSION

In this work, we extensively demonstrate the role of ANKRD26 in the regulation of three myeloid lineages by modulating the activity of three type I cytokine receptors that are essential in normal hematopoiesis.

We confirm a role for ANKRD26 in the regulation of the MK lineage, where its presence is crucial in the early steps of differentiation, but gets in the way of a correct terminal maturation. Indeed, shRNA-mediated decrease of ANKRD26 expression in CD34^+^ progenitors greatly reduces plating efficiency and the proliferative potential of MK progenitors. On the other hand, overexpression of ANKRD26 prevents the correct proplatelet formation in MKs, in agreement with the observations made for THC2 patients^8^.

For the first time, we show here that ANKRD26 plays a crucial role also in the granulocytic and erythroid lineages, the other two main myeloid lineages governed by a type I homodimeric receptors, G-CSFR and EPOR respectively. At the immature stage, hematopoietic stem and progenitors cells express basal levels of ANKRD26. Gene down-regulation in those cells lead to decreased proliferation and clonogenic potential, for the megakaryocytic, granulocytic and erythroid lineages, suggesting an important role of this protein in committed progenitors. Physiologically, ANKRD26 expression is progressively silenced along the cellular differentiation and maturation of those lineages. Our results show that an abnormal ANKRD26 expression in granulocytic lineage of THC2 patients impacts the normal process of cellular proliferation/differentiation that could lead to the leucocytosis reported in some patients. This mechanism could be responsible for the establishment of a fertile substrate prone to the acquisition of secondary mutations, justifying the increased incidence of myeloid malignancies detected in THC2 pedigrees ^6^.

Previously we have shown that the sustained ANKRD26 expression in the late phase of megakaryopoiesis lead to a defect in proplatelet formation, due to the deregulation of the TPO/MPL signaling.^8^ Here we demonstrate that the modulation of ANKRD26 expression levels modifies the TPO sensitivity. Interestingly, although ANKRD26 expression does not change the MPL levels at the cell surface, its higher expression increases the MPL signaling activity, leading to enhanced proliferation. This observation prompted us to hypothesize that ANKRD26 could be part of a complex that regulates the internalization of the receptor. By different approaches, we show that ANKRD26 and MPL are in the same protein complex, although further studies are needed to determine if their interaction is direct or not.

Finally, we show that both G-CSFR and EPOR interact with ANKRD26. Interestingly, thrombocytopenia rather than thrombocytosis is detected in TCH2 patients. This observation could be explained by the fact that, despite an amplification of MK progenitors due to the increased JAK2/STAT signaling at the early stage of MK-poiesis, the inactivation of MAPK-pathway necessary for correct proplatelet formation is not achieved^8^. In erythroid and granulocytic lineages, ANKRD26 modulates the proliferation rate, but not the late stages of differentiation. This feature could explain the reduced severity of the observed phenotypes for the erythroid and myeloid lineages. Moreover, the position of the pathogenic mutation could have a different impact on the binding of lineage-specific transcription factor(s), therefore regulating differentially the ANKRD26 expression level in the granulocytic and erythroid lineages.

Leukocytosis and erythrocytosis are often a consequence of mutations that alter the correct signaling of G-CSFR and EPOR. *CSF3R* mutations are recurrent in chronic neutrophilic leukemia (CNL), which is characterized by excessive hyperproliferation of neutrophils^37-39^, while *EPOR* mutations are a hallmark of familial erythrocytosis^27,40^. In both cases, the mutations often lead to the generation of truncated receptors that lack the specific domains that control the negative regulation of the signaling. We hypothesize that the persistent presence of ANKRD26 could act similarly to these mutations, by disturbing the receptors correct internalization and shutdown. Several reports link ANK repeat-containing proteins to the interaction and regulation of signaling pathways and receptor internalization. Ankrd26, the mouse homolog of ANKRD26, is involved in obesity onset^41-43^, via regulation of signaling pathways^44^, and ANKRD26 was also described to interact with different proteins, including hyaluronan-mediated motility receptor (HMMR)^45^. The ANK-domain containing protein ANKS1A participates in the endocytic trafficking of the EGFR by an unknown mechanism^46^. The endocytosis of EGFR is also regulated by ANKRD13A, 13B, and 13D, all containing an ubiquitin-interacting motif (UIM) through which they bind to the ligand-activated EGFR and direct its rapid internalization, probably by connecting the receptor with the endocytic machinery via their ANK domain^47^.

How ANKRD26 exactly interferes in the regulation of type I cytokine receptor internalization and signaling is not completely clear. ANKRD26 does not contain the UIM domain, and according to the recently reported thrombocytopenia caused by WAC-ANKRD26 fusion, with conserved C-but not N-terminal part containing ANK repeats^31^, only the C-terminal domain with coiled-coil region seems to be necessary for the interactions with MPL, EPOR and G-CSFR. Further studies are necessary to clarify if these interactions are direct or not and by which precise mechanisms ANKRD26 prevents their internalization.

In conclusion, we demonstrate a novel central role for ANKRD26 as responsible for the fine-tuning of at least three different receptors physiology. Small changes in their activity induce notable anomalies in proliferation of megakaryocytes, erythrocytes and granulocytes, and differentiation of megakaryocytes, which cause respectively thrombocytopenia, erythrocytosis and leukocytosis.

## ACKNOWLEDGEMENTS

The authors thank the patients for participation in this study, Pr. M.C. Alessi who coordinate the “Centre de Référence des pathologies plaquettaires” (France), P. Rameau, C. Catelain, T. Manoliu from the Imaging and cytometry platform PFIC, UMS AMMICA, Gustave Roussy Villejuif, France for the expertise in cytometry. M. K. Diop and T. Dayris from Bioinformatic platform, UMS AMMICA for RNA-seq analysis, Gustave Roussy Villejuif, France. M. Breckler from genomic platform UMS AMMICA, Gustave Roussy Villejuif, France for RNA-seq performed thanks to the TA2016_P27_ALDO. AP-HP, Hôpital Saint-Louis, Unité de Thérapie Cellulaire, CRB-Banque de Sang de Cordon, Paris, France for cord-blood samples. FBV, AD, VTM were supported by the Université Sorbonne Paris Cité/Université Paris Diderot and French Society of Hematology (FBV and VTM) and Ligue National Contre le Cancer (AD). The work of H.R. team is supported by grant from the Ligue Nationale Contre le Cancer (Equipe labellisée 2016 and 2019 to HR), and H2020-FETOPEN-1-2016-2017-SilkFusion.

## AUTHOR CONTRIBUTIONS

FBV and AD designed and performed experiments, analyzed data, contributed to the manuscript draft. VTM, ML, NB, CPMO, HD, BA, DM, EM and AC performed experiments and analyzed data. ND provided plasmid construction, supervised RNA-seq and discussed results. IB provided analysis of iPSC lines and discussed results. AB, ND, IAD, CM, IP, WV designed experiments, discussed results and contributed to manuscript editing. RF and LF provided patient and control samples and discussed results. HR designed and supervised the work and wrote the paper. FBV and AD equally contributed to this work. All the authors contributed to the final approval of the manuscript.

