## Supplementary Tables, Supplementary Figures, Supplementary Material and Methods for "ANKRD26 is a new regulator of type I cytokine receptor signaling in normal and pathological hematopoiesis"

**Running title:** ANKRD26 regulates cytokine-mediated signaling

### MATERIAL AND METHODS

#### Study approval

Peripheral blood samples from patients and healthy subjects, and cord-blood samples were collected after informed written consent and obtained in accordance with the Declaration of Helsinki. The study was approved by the Comité de Protection des Personnes CPP N°2020T2-02 and by AP-HP, Hôpital Saint-Louis, Unité de Thérapie Cellulaire, CRB-Banque de Sang de Cordon, Paris, France, N° of authorization: AC-2016-2759.

The NSG mice were provided from Charles River Laboratories. Animal experiments were performed in accordance with 2010/63/UE European legislation and decree n°2013-118 of French legislation and recorded under protocol number APAFIS# 2016-008-7175. Mice were housed in pre-clinical platform of Gustave Roussy Animal Facilities PFEP (Villejuif, France) (ministerial approval n° 94-076-11).

#### Plasmid constructions

The full cDNA of human ANKRD26 was amplified by PCR from the pFN21a_ANKRD26_Halotag plasmid, using Phusion High-Fidelity Polymerase (Thermo Scientific) and ANKRD26_pEF6/V5-His primers. The PCR product was digested with DpnI enzyme (Thermo Scientific) for 1h at 37°C, to remove the wild-type template plasmid, purified using GeneJET Gel Extraction Kit (Thermo Fisher) and cloned into the linearized pEF6/V5-His TOPO vector using the InFusion kit (Clontech), following the manufacturer's instructions. Then, the ANKRD26_V5-His sequence was amplified by ANKRD26_pRRL primers and cloned into the linearized pRRL_EF1a-MCS-PGK_hygromycin B or pRRL_EF1a-MCS-PGK_Cherry lentiviral vector. Alternatively, ANKRD26 cDNA was amplified from the pFN21a_ANKRD26_Halotag plasmid using FLAG-ANKRD26 primers and cloned into the linearized pRRL_EF1a-MCS-PGK_hygromycin B lentiviral vector. The HA_G-CSFR cDNA was amplified by RT-PCR with primers containing HA sequence and cloned into the MIGR retroviral plasmid. Plasmids were obtained after transformation of XL_10 Gold ultracompetent cells and DNA purification with (NucleoBond Xtra Midi kit, Macherey-Nagel), and constructions were verified by sequencing. Lentiviral vectors encoding either for short hairpin RNAs against ANKRD26 (sinPRRL-H1-shANK-PGK-mCherry or sinPRRL-H1-shANK-PGK-GFP) or a scramble sequence (sinPRRL-H1-shSCR-PGK-mCherry or sinPRRL-H1-shSCR-PGK-GFP) were previously described[8]. Two different shANK (shANK_1 and shANK_2)[8] giving similar results were used and the results are presented as a mean of both. A retrovirus encoding for wild type Jak2 was used (pREX- Jak2_IRES-CD4) was used for PLA assay. Two different shRNA against CCNI2 (shCCNI2_1 and shCCNI2_2 were cloned) into the sinPRRL-H1-PGK-GFP vector (sinPRRL-H1-_CCNI2_PGK-GFP).

**Lenti- and Retroviral production**

For lentiviral production, HEK293T cells were seeded to achieve 70% confluence at the time of transfection, typically $1x10^6$ cells per T175 flask seeded three days before. The cells were co-transfected with the following plasmids: pCMV, pMD2G, and the plasmid of interest. For retroviral production, HEK293EBNA cells were co-transfected with the following plasmids: GAG-POL, VSV-G, and the plasmid of interest. Transfection was carried out using JetPrime reagents following the manufacturer's instructions. The viral supernatant was collected 48 and 72 hours after transfection, concentrated by ultracentrifugation at 22K rpm for 1h30, and stored at -80°C.

**Primary cell culture**

$CD34^+$ cells were isolated from umbilical cord blood or peripheral blood by positive selection using immunomagnetic bead cell-sorting system (AutoMacs; Miltenyi Biotec) and cultured in Iscove modified Dulbecco medium with penicillin/streptomycin/glutamine, alpha-thioglycerol, bovine serum albumin, a mixture of sonicated lipids and insulin-transferrin containing SCF (25 ng/mL) and TPO (10 ng/mL), or G-CSF (20 ng/mL), IL-3 (10 ng/mL) and SCF (25 ng/mL) or apo-transferrin instead of insulin-transferrin, EPO (1 U), IL-3 (10 ng/mL) and SCF (25 ng/mL).

**Cell lines**

**Human UT7** cells were maintained in MEMα medium supplemented with 10% fetal calf serum, 1% PenStrep, 1% L-Glutamine, and in the presence of 5 ng/mL of GM-CSF. They were transduced with lentiviral vectors encoding either for short hairpin RNAs against ANKRD26 or a scramble sequence, as previously described[8], and isolated via cell sorting 48 hours later, based on the reporter gene expression. Similarly, UT7 cells were transduced with a lentiviral vector encoding for the full-length ANKRD26 cDNA expressed under the EF1a promoter and a hygromycin B resistance gene expressed under PGK promoter (pRRL-EF1a-ANKRD26/PGK-hygromycin B) and were selected using 400 µg/mL of antibiotic.

**Murine Ba/F3** cells were maintained in DMEM medium supplemented with 10% fetal calf serum, 5% Wehi, 1% PenStrep, 1% L-Glutamine, and in the presence of 5% WEHI-conditioned medium. Ba/F3 were transduced with the retroviral vector encoding for wild-type human HA_MPL, HA_EPOR or HA_G_CSFR and the lentiviral vector pRRL_EF1a-ANKRD26/PGK-hygromycin B or the empty vector pRRL_EF1a-MCS_PGK-hygromycin B. Human Gamma-2A cell line without expression of endogenous JAK2 and Human embryonic kidney HEK293T cell lines were maintained in DMEM medium supplemented with 10% fetal calf serum, 1% PenStrep, 1% L-Glutamine.

All cells were cultured in a humidified incubator at 37°C with 5% CO2.

**iPSCs expansion**

iPSCs were maintained on Essential 8 or Essential 8 Flex medium (Gibco), on plates coated with N-truncated Human recombinant vitronectin (Gibco). Cell passage was routinely performed using a solution of EDTA 500 M in PBS 1X, or TrypLE 1X (Gibco). A Mycoplasma screening was routinely performed, accordingly to the manufacturer instructions (Sigma). Cells were kept in culture for a limited number of passages, to prevent the surge of karyotype and genomic anomalies. The characterization of iPSCs is described in supplemental material and method section.

**iPSC characterization**

***qRT-PCR analysis of pluripotency markers***

The expression of self-renewal stem cell markers was validated by qRT-PCR analysis. qRT-PCR was assessed by the DDCt method on Rotor-Gene Q (Qiagen) using the Maxima SYBR Green qPCR Master Mix (ThermoFisher Scientific). GAPDH was selected as a housekeeping gene and data were normalized to its expression. Statistical analysis was performed using REST (Relative Expression Software Tool) software. The expression of self-renewal stem cell markers of ANK1 and ANK2 cells was compared to hESCs RC17 (Rosalin Cells, Edinburg UK) and hiPSCs CTR2#6 and C1[28].

***Immunofluorescence analysis of pluripotency markers***

The expression of self-renewal stem cell markers was validated by immunofluorescence analysis performed on cells 48h after thawing. Cells were immunostained at 4 °C overnight with primary antibodies, and with appropriate secondary antibodies for 1 hour at RT; nuclei were counterstained with Hoechst 33258. The images were acquired using a Leica DMI4000B inverted microscope linked to a DFC360FX or to a DFC280 camera (Leica Microsystems).

***Karyotype Analysis***

iPS-cells were treated on the 4th day after split for 3 hours with 0.1 μg/ml Colcemid. A total of 32 to 38 metaphases were analyzed and 13-15 total metaphases were karyotyped by QFQ-banding: q-Bands by Fluorescence using Quinacrine and with Fluorescence Microscope Olympus CHB (BX63) with quinacrine mustard filter and CCD camera Bright-Field Microscope"GenASIs" Software, version 8.1.0.47741; Applied Spectral Imaging.

***Short Tandem Repeat (STR) Testing Report***

The amplification was performed by Polymerase chain reaction (PCR) for each nine autosomal STR molecular markers (D21S11, D7S820, CSF1PO, TH01, D13S317, D16S539, vWA, TPOX, D5S818) along with the gender-determining marker Amelogenin with Promega GenePrint 10 Kit following manufacturer's recommended protocol. Appropriate positive and negative amplification controls were used as kit recommended guidelines. The amplified products were analyzed on an ABI Prism® 3730xl Genetic Analyzer using an Internal Lane Standard 600 (Promega). Data generated was analyzed using GeneMapper® Software version 4.0 (Applied Biosystems) following manufacturer's instructions.

***Teratoma formation assay***

iPS-cells ($1x10^6$ cells) were resuspended in Essential 8 (140 L) and Geltrex (60 L) to a final volume of 200 L and injected intramuscularly in NSG mice. Mice were monitored and euthanized when the size of the tumoral mass was visibly affecting the animal motility and behavior. Teratomas were excised, fixed and embedded in paraffin, while the corresponding sections were stained with hematoxylin, eosin and safranin.

**iPSCs hematopoietic differentiation**

Clumps of pluripotent cells were seeded on Geltrex (Gibco)-coated plates, in presence of E8 medium, at day -1. The departing cell concentration was adjusted for each cell line and was comprised in the 10-20% confluence range. At Day 0, cells were transferred in a xeno-free medium based on StemPro-34 SFM (Gibco), supplemented with Penicillin/Streptomycin 1% v/v (Gibco), L-Glutamine 1% v/v (Gibco), 1-Thioglycerol 0.04 mg/mL (Sigma) and ascorbic acid 50 μg/mL (Sigma). This medium was retained for the entire experiment and supplemented with different cytokines and growth factors, accordingly to the following schedule: Days 0 – 2:

BMP4 (10 ng/mL), VEGF (50 ng/mL) and CHIR99021 (2 M). Days 2-4: BMP4 (10 ng/mL), VEGF (50 ng/mL) and FGF2 (20 ng/mL). Days 4 – 6: VEGF (15 ng/mL) and FGF2 (5 ng/mL). Day 6: VEGF (50 ng/mL), FGF2 (50 ng/mL), SCF (50 ng/mL) and FLT3L (5 ng/mL). Days 7-10: VEGF (50 ng/mL), FGF2 (50 ng/mL), SCF (50 ng/mL), FLT3L (5 ng/mL), TPO (50 ng/mL) and IL-6 (10 ng/mL). Days 10-20: SCF (50 ng/mL), G-CSF (25 ng/mL), and IL-3 (10 ng/mL).

**Clonogenic potential of primary cells in semi-solid culture**

***Methylcellulose culture assay:*** hematopoietic progenitors ($CD34^+CD43^+$ for iPSC and $CD34^+$ for primary cells) were plated in triplicate at different cell densities ($1x10^3$ cells/plate for primary cells, $3x10^3$ cells/plate for iPSC-derived cells) in human methylcellulose medium H4434 (STEMCELL technologies), containing recombinant human cytokines and scored for the presence of colonies 14 days after. When indicated, $CD34^+$ progenitor cells transduced with lentiviral vectors encoding for short hairpin RNAs against ANKRD26 (PRRL_H1-shANK/PGK-GFP) or a scramble sequence (PRRL_H1-shSCR/PGK-GFP), were cultured for 2 days in presence of SCF (25 ng/mL), FLT3L (10 ng/mL), TPO (10 ng/mL), IL-6 (10 ng/mL), IL-3 (10 ng/mL). $CD34^+GFP^+$ cells were then sorted on a BD Influx instrument and seeded on triplicate at $1x10^3$ cells/mL in methylcellulose H4230 (STEMCELL technologies) supplemented with 25 ng/mL SCF alone or with different doses of EPO (1, 0.1, 0.01 U/mL).

***Fibrin clot assay:*** To assess the CFU-MK potential, $CD34^+GFP^+$ or $CD34^+Cherry^+$ sorted cells were seeded at 1.5 x $10^3$ cells/mL in triplicate in fibrin clot medium, as previously described[29]. Different concentrations of TPO (0.01 ng/mL, 0.1 ng/mL, 1 ng/mL, 10 ng/mL) were added to 25 ng/mL SCF; plates with only SCF and no cytokines were also prepared. CFU-MK colonies were scored after 10 days, using an indirect alkaline phosphatase–based immunostaining technique based on CD41a detection by monoclonal antibody.

**Proliferation assay in liquid medium**

UT7/HA_MPL shANK/shSCR cells were washed three times in phosphate-buffered saline (PBS 1X) and seeded in MEMα with the indicated concentrations of TPO or Eltrombopag and 5 ng/mL of GM-CSF as control. $5x10^4$ cells per condition were plated in triplicate and manually assessed every day for 4 days, performing a Trypan blue exclusion and using a Malassez hemocytometer. The same protocol was used to measure the proliferation of UT7/HA_G-CSFR shANK/shSCR cells and UT7/HA_EPOR shANK/shSCR cells, using the appropriate cytokine and GM-CSF as positive control.

Primary cells ($CD34^+$) or iPS-derived progenitors ($CD34^+CD43^+$ sorted at day 14 of culture) were seeded at a cell density of $5x10^4$ cells/well in a 12-wells plate, in serum-free medium containing G-CSF (25 ng/mL), SCF (20 ng/mL) and IL-3 (10 ng/mL) for granulocytic lineage and EPO (1U/mL), SCF (20 ng/mL) and IL-3 (10 ng/mL) for erythroid lineage. Alternatively, iPS-derived $CD34^+CD43^+$ progenitors were sorted at day 14 of culture, transduced with shSCR or shCCNI2 encoding lentiviruses, sorted at day 16 on $GFP^+$ and $1x10^4$ cells/well were seeded in a 48-well plate. Cell numbers were quantified at the indicated days. The proliferation curves are reported relative to the number of seeded cells.

**MGG staining**

Cells were centrifuged on slides for 5 minutes at 500 rpm, on a Cytospin 2 centrifuge (Shandon). Slides were stained with May-Grünwald Giemsa (MGG); at least 200 cells per slide were scored. Images were obtained using a Leica DMRB microscope, using a 63x magnification objective. The % of myeloblasts, promyelocytes, myelocytes, metamyelocytes and PNN (based on the morphology) was calculated after counting under the microscope 200 cells corresponding to all these five subtypes.

**Proplatelet formation assay:** $CD41^+CD42^+GFP^+$ or $CD41^+CD42^+$ cells selected on hygromycin B were sorted at day 10 of culture and seeded at a cell density of $5x10^3$ cells/well in a 96-wells plate, in a serum-free medium containing TPO (10 ng/mL) and SCF (25 ng/mL). Proplatelet-forming cells were scored after 4-5 days by enumeration of no less than 200 cells per well using an inverted microscope (Carl Zeiss), at a 200x magnification. A proplatelet-forming megakaryocyte was considered as a cell displaying at least one cytoplasmic process with a clearly defined constriction area. Each condition was examined in triplicate.

**Receptor surface expression and internalization assay**

UT7 cells expressing respectively HA_MPL, HA_G-CSFR or HA_EPOR and transduced either with shSCR or shANK were stained with an anti-MPL or anti-HA (Miltenyi) antibody and analyzed on a LSRFortessa flow cytometer (BD).

For internalization assay, Ba/F3-HA_MPL, Ba/F3-HA_MPL-ANKRD26_V5, Ba/F3-HA_EPOR, Ba/F3-HA_EPOR-ANKRD26_V5, Ba/F3-HA_G_CSFR and Ba/F3-HA_G_CSFR-ANKRD26_V5 cells ($2x10^5$ cells per time point) were washed twice in PBS 1x and cytokine-deprived overnight in IMDM medium supplemented with 1,5% bovine serum albumin (BSA). The following days, cells were stimulated with 50 ng/mL TPO, 1 U/mL RPO or 20 ng/mL G-CSF for 15/30/60 min, a non-stimulated condition was used as T=0. At the end of the stimulation, cells were washed in cold PBS supplemented with 0,5% BSA, incubated with a PE-conjugated anti-CD110 antibody (MPL-PE), an APC-conjugated anti-HA antibody (HA-APC) or a matched IgG for 30 min on ice, and then analyzed on a Canto X flow cytometer (BD). The median fluorescence intensity (MFI) was analyzed.

**Flow Cytometry**

Single cell suspensions were stained with monoclonal antibodies directly coupled to their respective fluorochromes. A list of the used antibodies, included the isotype control, is available, with the respective clone names (Supplementary Table 4). Cells were incubated with antibodies at the concentration of 1 L/$10^6$ cells, in approximately 100 L, at 4 °C for at least 30 minutes. Cells were washed before and after incubation in PBS 1X and analyzed with a BD Canto II or BD LSRFortessa cytometers (BD). Cell Sorting was routinely used for hematopoietic cells purification and performed on Influx, ARIA III or ARIA Fusion cell sorters (BD).

**Western blot analysis**

Signaling studies were performed on UT7 cell lines and iPS-derived cells

***UT7 cells*** expressing respectively HA_MPL, HA_G-CSFR and HA_EPOR and transduced with shSCR or shANK were washed three times in PBS 1X and cytokine-deprived overnight in MEMα medium supplemented with 5% FCS.

***Ba/F3 cells*** expressing respectively HA_MPL, HA_G-CSFR and HA_EPOR and transduced with pRRL_EF1a-MCS-PGK_hygromycin B (empty vector, EV) or pRRL_EF1a-ANKRD26-PGK_hygromycin B (selected by 400 μg/mL Hygromycin B) were washed three times in PBS 1X and cytokine-deprived overnight in IMDM medium supplemented with 1% BSA.

Re-stimulation was done with TPO/G-CSF/EPO for the indicated times and doses at 37°C. Next, cells were washed in ice-cold PBS to block stimulation. Total cell lysates were obtained by addition of 2X Laemmli buffer supplemented with DTT 0.1 M.

***iPS-derived cells:*** cells were washed three times in PBS 1X and starved overnight in serum-free medium, deprived of insulin-transferrin-selenium. At least $4x10^5$ cells per condition were plated. Re-stimulation was done with 25 ng/mL of G-CSF at different time points. Cells were then harvested in ice-cold PBS 1X, washed three times in clod PBS 1X, immediately lysed in ice-cold Laemmli buffer containing DTT 1 mM, protease inhibitor cOmplete (5X)

(cOmplete™, Mini Protease Inhibitor Cocktail, Roche), phosphatase inhibitor cocktail 3 (25X), sodium orthovanadate 50 mM, sodium fluoride 500 mM and PMSF 100 mM.
Lysates were then sonicated and heated at 95°C for 5 minutes. Proteins were resolved by SDS-PAGE and then transferred onto a nitrocellulose membrane. After blocking in 5% BSA in TBS-0.1% Tween20 (TBS-T) for 1 hour, the membranes were incubated with rabbit primary antibodies (anti–phospho-STAT5 (Y705), anti-STAT5, anti–phospho-STAT3 (Y705), anti-STAT3, anti–phospho-AKT (S473), anti-AKT, anti–phospho-ERK1/2, anti-ERK1/2 (Cell Signaling Technologies), diluted 1:1000 in blocking buffer overnight at 4°C with gentle shaking. For detection, the membranes were washed and incubated with secondary HRP-conjugated antibodies, diluted 1:5000 in blocking buffer. Band detection was performed by enhanced chemiluminescence system (ECL or SuperSignal West Pico Plus kit; Life Technologies) using Image Quant LAS 4000 (GE Healthcare)/iBright FL1500 (Thermo Fisher Scientific). The quantification was by ImageJ2 software.

**Co-immunoprecipitation**

HEK293T cells were transiently transfected with a pMX-IRES-GFP vector encoding respectively for HA_MPL, HA_EPOR and HA_G-CSFR, and with a pEF6/V5-His TOPO vector encoding for ANKRD26, using JetPrime reagents. After 72 hours, 1-2 x $10^7$ cells were pelleted by centrifugation and lysed in a buffer containing 50 mM TrisHCl pH 7.5, 150 mM NaCl, 1% NP40, supplemented with fresh protease inhibitors (cOmplete™, Mini Protease Inhibitor Cocktail, Roche) and phosphatase inhibitors (1 mM Na3VO4, 10 mM NaF). Lysates were cleared by centrifugation at 10,000 g for 15 min at 4°C and then incubated with primary antibodies (anti-TPOR 06-944, Millipore; anti-V5 R960-25, Invitrogen; anti-HA ab9110, Abcam) overnight at 4°C with gentle shaking. As no specific antibodies are available for EPOR and G-CSFR, anti-HA was used in these two cases. Next, protein A/G Plus agarose beads (sc-2003, Santa Cruz Biotechnology) were added for 2 hours at 4 °C. Beads were washed once with lysis buffer and once with PBS, and samples were boiled in 2X Laemmli sample buffer supplemented with DTT 0.1 M. Samples were then analyzed by western blotting as described above.

**Proximity ligation assay (PLA)**

Parental UT7 cells were transduced with FLAG_ANKRD26_Hygromycin and HA_MPL_GFP encoding lenti- or retroviruses, sorted on $GFP^+$ and cultured in presence of 400 µg/mL Hygromycin B (Sigma Aldrich). Non-transduced cells were used as negative controls.

Alternatively, adherent Gamma-2A cells were transiently transfected with FLAG_ANKRD26_Hygromycin, HA_MPL_GFP and Jak2_CD4 lenti- or retroviruses or corresponding empty vectors by using TransIT®-293 transfection reagent (Mirus). First, the cells were transfected with FLAG_ANKRD26_Hygromycin lentivirus or corresponding empty vector and cultured in presence of 400 µg/mL of Hygromycin B (Sigma Aldrich). Then, cells were transfected with HA_MPL_GFP and/or Jak2_CD4 retroviruses. Two days after second transfection, cells were stained with anti-CD4-APC antibody and $CD4^+GFP^+$ cells were sorted. After sorting, UT7 or Gamma-2A cells were cytocentrifuged (5 min at 500 rpm) to Poly-L-Lysine slides (Thermo Fisher Scientific). Cells were fixed with 4% paraformaldehyde in for 10 min, then permeabilized in PBS 1x, containing 0.2% Triton X-100 for 5 minutes. Fixed cells were incubated overnight with anti-FLAG (F7425, rabbit, MilliporeSigma, dilution 1:400) and anti-HA (901513, mouse, BioLegend, dilution 1:200) antibodies. Proximity ligation assays were performed according to the manufacturer's instructions (Duolink, Sigma Aldrich) using oligonucleotide-coupled secondary antibodies against mouse and rabbit primary antibodies (PLA probes). Only when a pair of PLA probes has bound two primary antibodies in close proximity (<40 nm), a red fluorescent spot is generated as a result of the hybridization and circularization of fluorescently labeled oligonucleotides during the amplification reaction. No fluorescence was detected when the cells were incubated in the presence of the two complementary oligonucleotides and the kit components, but without primary antibodies. Images were acquired under a Confocal Leica SP8 laser-scanning microscope, with a 63x/1.4 numeric aperture oil objective (Leica Microsystem). Slides analyzed for a presence of red dots were scored by enumeration of at least 100 cells per slide. Image analysis was performed with the LASX software. For each experiment at least 10 cells per condition (for UT7) and 40 cells per condition (for Gamma-2A) were analyzed for the number of spots using Image J2 software.

**Whole-transcriptome RNA-seq and analysis**

Cells were sorted and flash-frozen at -80 °C. RNA extraction and purification were performed with the RNeasy Mini Kit (Qiagen). The RNA integrity (RNA Integrity Score≥7.0) was checked on the Agilent 2100 Bioanalyzer (Agilent) and quantity was determined using Qubit

(Invitrogen). SureSelect Automated Strand Specific RNA Library Preparation Kit was used according to manufacturer's instructions with the Bravo Platform. Briefly, 50 to 200ng of total RNA sample was used for poly-A mRNA selection using oligo(dT) beads and subjected to thermal mRNA fragmentation. The fragmented mRNA samples were subjected to cDNA synthesis and were further converted into double stranded DNA using the reagents supplied in the kit, and the resulting dsDNA was used for library preparation. The final libraries were bar-coded, purified, pooled together in equal concentration and subjected to paired-end sequencing on Novaseq-6000 sequencer (Illumina).

Quality controls were assessed on raw fastq with RseqQC and FastQC. Remaining rRNA were filtered out with BWA, on UCSC's HG19. Filtered reads trimming was performed with Trimmomatic using official TrueSeq adapter list, filtering qualities lower than 20 on a sliding window of size 6, and a minimum length of 40 bases. Remaining reads were mapped with TopHat2 against UCSU's HG19. The insert length deducted in previous RSeQC step, and the realignment edition distance parameter set to 0 were used as optional parameters. Counts were performed over Samtool-sorted bam with HTSeqCount. Rows full of zeros and non-genic annotations from HTSeqCount were removed through in-house bash scripts. The differential analysis was performed with DEseq2, using VST normalization method and default GLM model. Quality control graphs and results were performed with in-house R-scripts (available on request). The data were submitted to EGA under the number EGAD00001008023.

GSEA was performed using the software GSEA (v4.1.0).

**Quantitative Real Time Polymerase Chain Reaction (qRT-PCR)**

RNA was purified using the TRIzol™ Reagent (Invitrogen), RNeasy Plus Micro or Mini Kit, accordingly to the cell number and the manufacturer instructions (Qiagen). Complementary DNA synthesis was performed with the Super Script II cDNA synthesis Kit or the Super Script Vilo cDNA synthesis kit (Thermo Fisher), accordingly to the amount of extracted RNA. Real Time PCRs were performed using the Takara Bio SYBR Premix Ex Taq (Clontech) reaction mix, on the 7500 Real Time PCR system (Applied Biosystems). Gene expression was assessed by comparative CT method, using PPIA or HPRT as reference genes.

### Supplementary Figures

#### Supplementary Figure 1

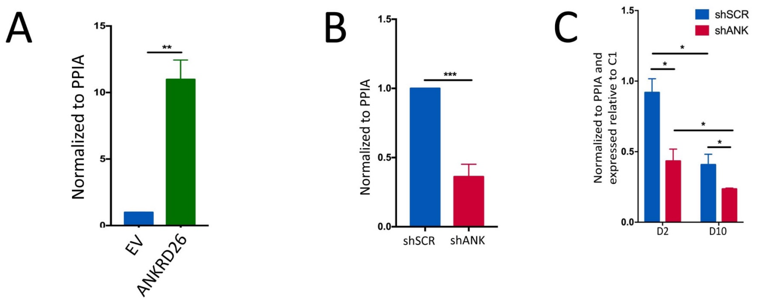


**Supplementary Figure 1. Quantification of ANKRD26 transcripts.**
A) *ANKRD26* transcript level in cord-blood (CB)-CD34$^+$ cells transduced with empty lentivirus (EV) or lentivirus encoding ANKRD26_V5 and Cherry. CD34$^+$Cherry$^+$ cells were sorted at day 2 post-transduction. *ANKRD26* transcript level was measured by qRT-PCR and normalized to *PPIA* housekeeping gene. B) CB-CD34$^+$ hematopoietic progenitors were transduced with lentiviruses encoding shSCR (control) or shANK (shANK_1 or shANK_2) and GFP. CD34$^+$GFP$^+$ cells were sorted at day 2 post-transduction. C) CB-CD34$^+$ hematopoietic progenitors were transduced with lentiviruses encoding shSCR (control) or shANK (shANK_1 or shANK_2) and GFP. *ANKRD26* transcript level was measured in CD34$^+$GFP$^+$ cells sorted at day 2 post-transduction (D2) and in megakaryocytes at day 10 (D10) of culture in presence of TPO and SCF.
*ANKRD26* level was measured by qRT-PCR and normalized to *PPIA* housekeeping gene. Shown are averages of 3 independent experiments as mean±SD. In A and B, the results are presented relative to EV. *P<0.05; **P<0.01; ***P<0.005, paired t-test.

### Supplementary Figure 2

A

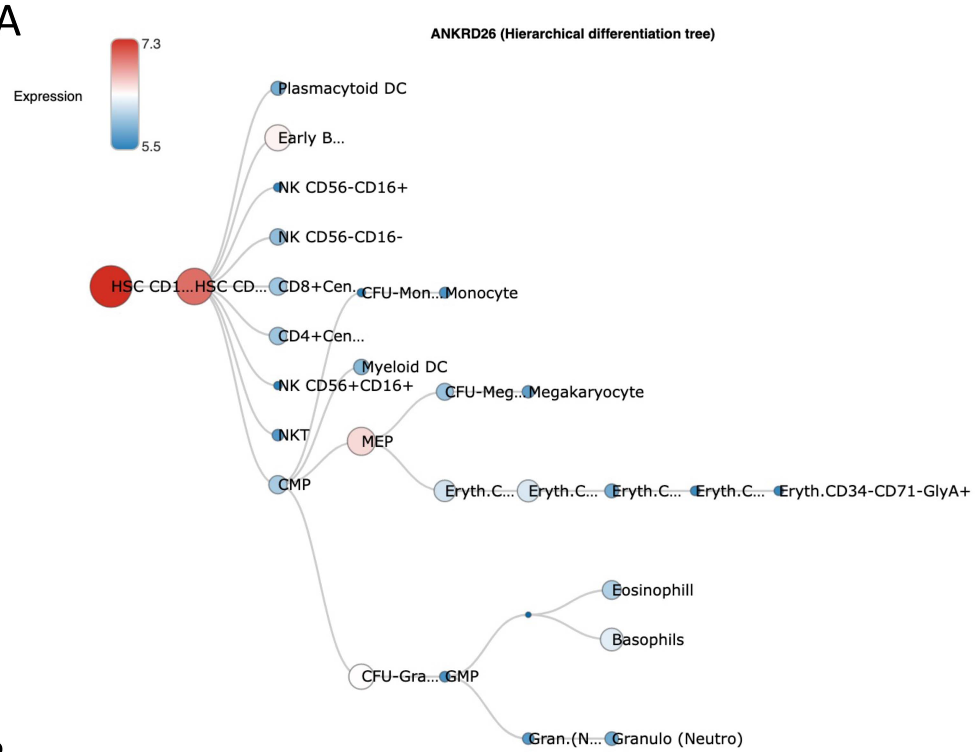


B

From: https://servers.binf.ku.dk/bloodspot/?gene=ANKRD26&dataset=DMAP

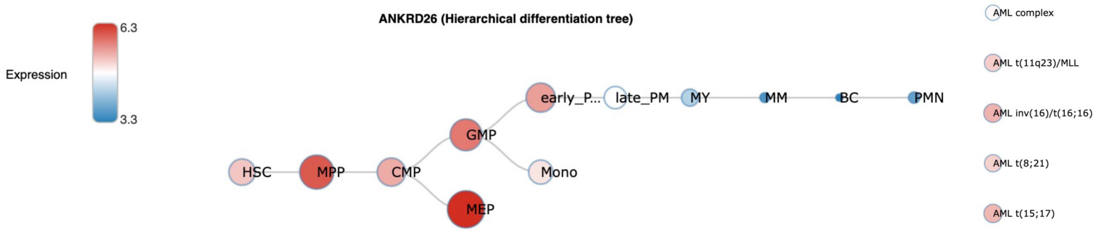


From: https://servers.binf.ku.dk/bloodspot/?gene=ANKRD26&dataset=normal_human_v2_with_AMLs

**Supplementary Figure 2. ANKRD26 expression level through hematopoietic differentiation.**

The data are extracted from BloodSpot database (https://servers.binf.ku.dk/bloodspot/). ANKRD26 is highly expressed in hematopoietic stem cells (HSC) and multipotent progenitors (MPP) (A, B), and its expression level decreased through megakaryocyte (A), erythrocyte (A) and granulocyte (A, B) differentiation. ANKRD26 is expressed in different AML subtypes (B).

### Supplementary Figure 3

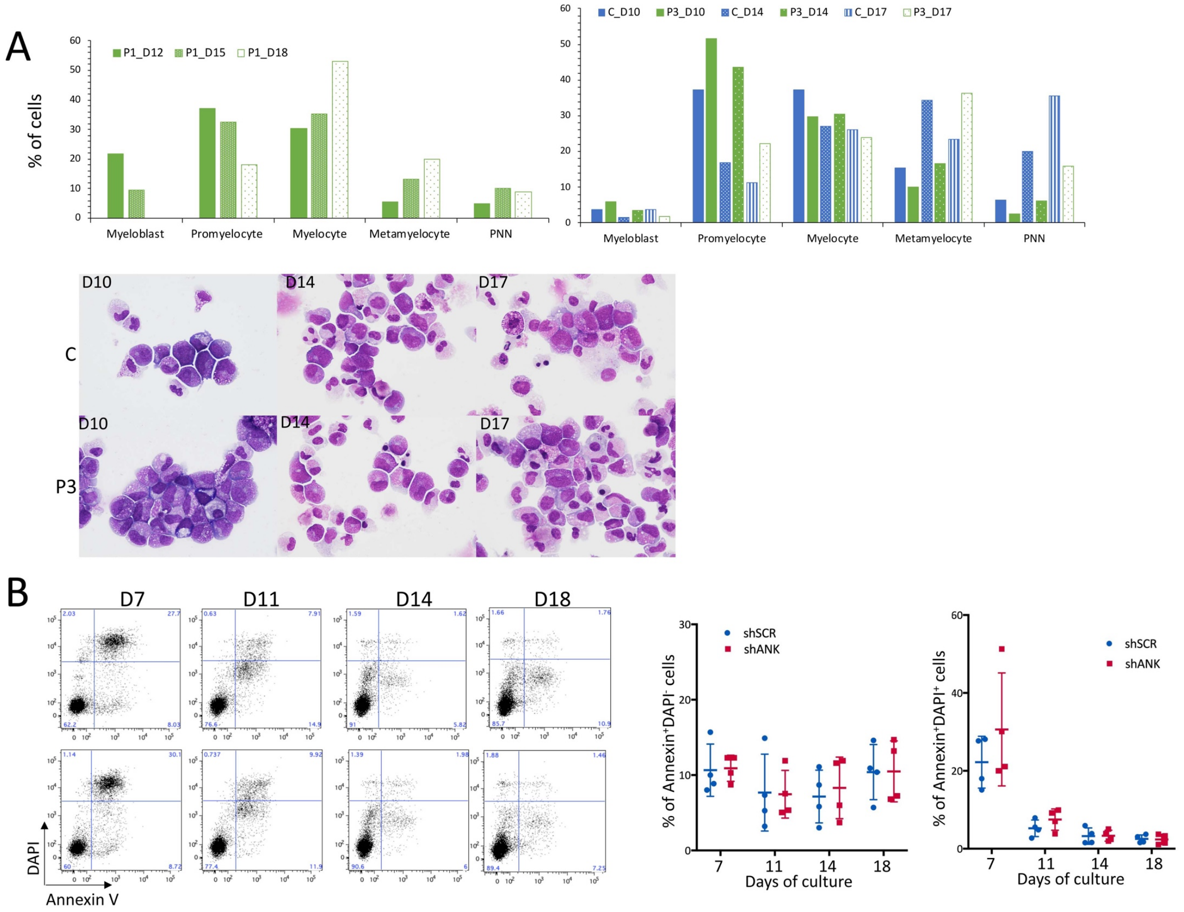


**Supplementary Figure 3. ANKRD26 levels affect granulocyte maturation, but not apoptosis.**

A) Kinetics of granulocyte differentiation for one control and two patient samples. The figure shows that ANKRD26 overexpression induces a delay, but not a blockage of granulocytic differentiation *in vitro* in THC2 patients. MGG staining and analysis of the proportion of myeloblasts, promyelocytes, myelocytes, metamyelocytes and PNN was performed at different days of culture of $CD34^+$ cells in presence of G-CSF, IL3 and SCF. MGG staining corresponding to 1 control and 1 patient sample is shown in the lower panel. B) Effect of ANKRD26 inhibition on apoptosis in granulocytic lineage. $CD34^+$ cells were transduced with lentiviruses encoding respectively shSCR and shANK (shANK_1/shANK_2), and were sorted 2 days after transduction. They were grown in liquid medium in presence of 25 ng/mL SCF, 10 ng/mL IL-3 and 20 ng/mL G-CSF. Apoptosis (Annexin-$V^+DAPI^-$ and Annexin-$V^+DAPI^+$) was measured at different days of culture. Shown are the averages of 3 independent experiments as mean±SD.

**Supplementary Figure 4.**

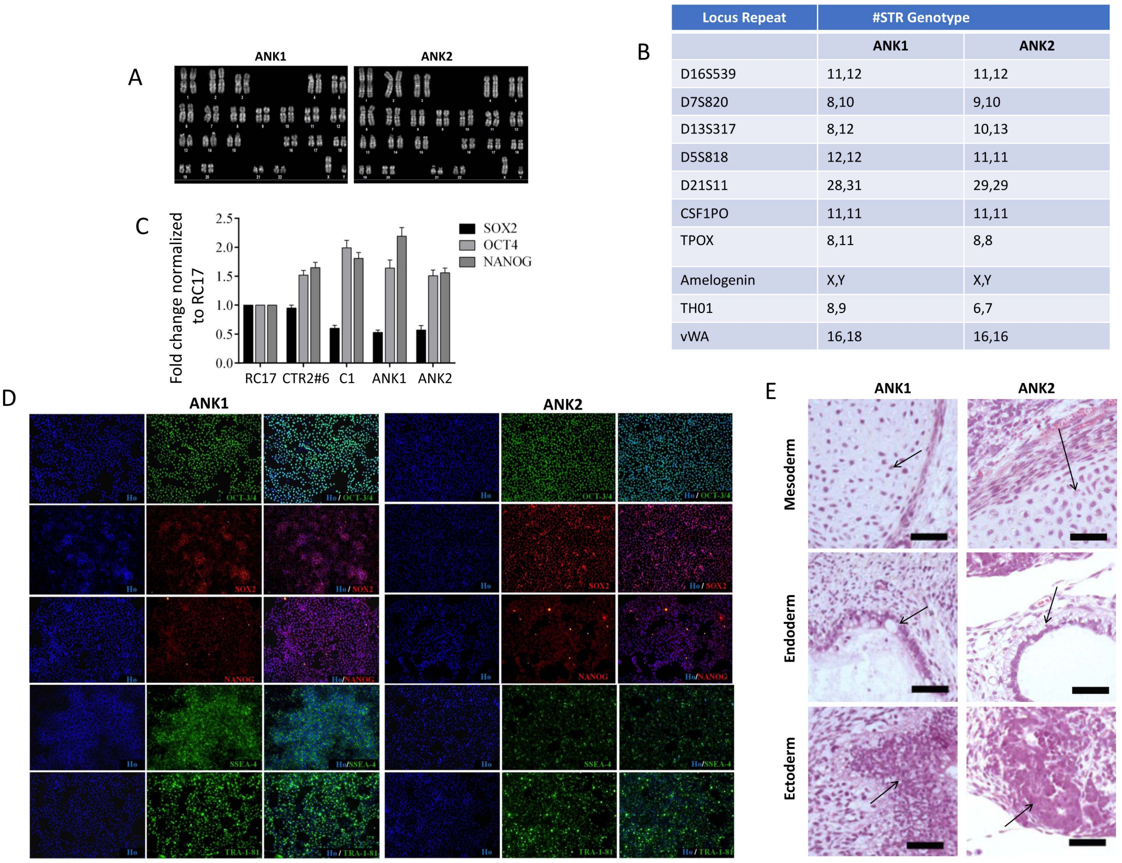


| Locus Repeat | #STR Genotype | |
|---|---|---|
| | ANK1 | ANK2 |
| D16S539 | 11,12 | 11,12 |
| D7S820 | 8,10 | 9,10 |
| D13S317 | 8,12 | 10,13 |
| D5S818 | 12,12 | 11,11 |
| D21S11 | 28,31 | 29,29 |
| CSF1PO | 11,11 | 11,11 |
| TPOX | 8,11 | 8,8 |
| Amelogenin | X,Y | X,Y |
| TH01 | 8,9 | 6,7 |
| vWA | 16,18 | 16,16 |

**Supplementary Figure 4. Characterization of iPSC ANK1 and ANK2 lines.** A) Karyotype analysis showing normal 46, XY karyotype for both ANK1 and ANK2 iPSC lines. 32 to 38 total metaphases were analyzed and 13-15 total metaphases were karyotyped by Q-binding method with resolution of 350 bands. B) Short Tandem Repeat (STR) Testing Report. C-D) Analysis of pluripotency markers. C) qRT-PCR analysis of *SOX2*, *OCT4* and *NANOG* transcripts relative to *GAPDH* housekeeping gene and normalized to control line hESCs RC17 cell line. D) Immunofluorescence analysis of pluripotency markers OCT3/4, SOX2, NANOG, SSEA-4, TRA-1-81. E) *In vivo* differentiation of iPSC ANK1 and ANK2 lines into 3 germ layers; mesoderm, endoderm and ectoderm.

**Supplementary Figure 5**

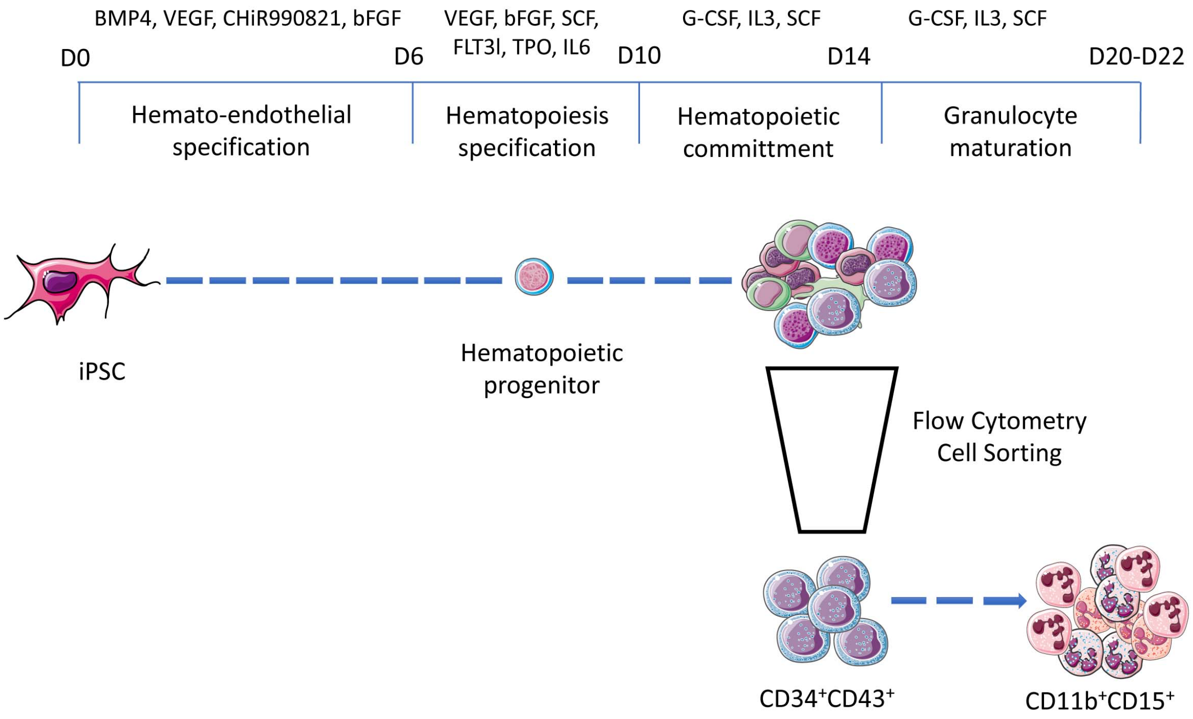


**Supplementary Figure 5**. **Schema of granulocyte differentiation protocol.** D indicates days of culture.

**Supplementary Figure 6**

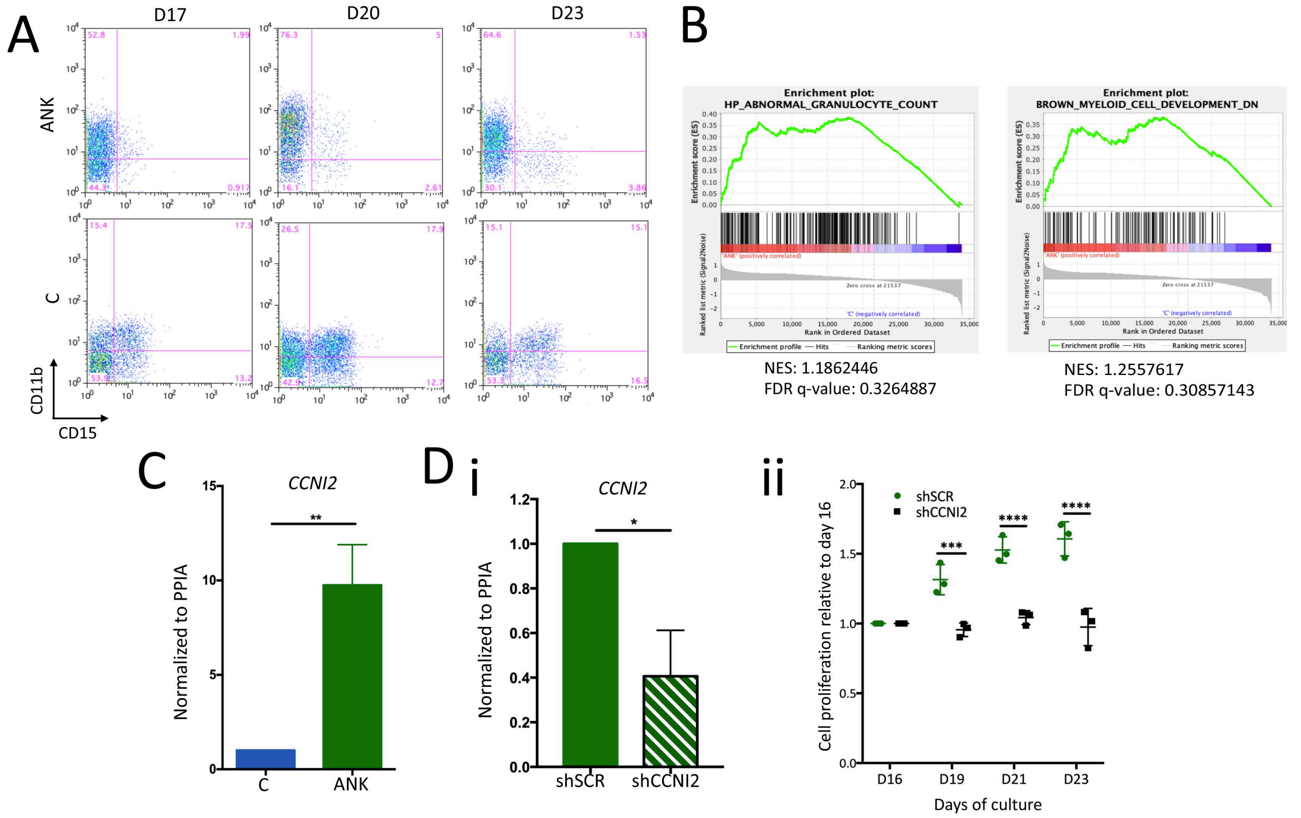


**Supplementary Figure 6. Increased ANKRD26 expression level enhances proliferation of granulocyte progenitors in an iPSC model.**

A) FACS analysis of $CD34^{+}CD43^{+}$ progenitors derived from patient (ANK) and control (CTR) iPSC differentiated as granulocytes in the presence of G-CSF, SCF and IL-3 at different days of culture. B) RNA-seq assay was performed on $CD43^{+}CD11b^{+}CD14^{-}$ progenitors sorted at day 16 of culture. Shown are GSEA analysis for HP_ABNORMAL_GRANUCOLYTE_COUNT (HP:0032309) and for BROWN_MYELOID_CELL_DEVELOPMENT_DN in ANK versus Control iPSC. C) qRT-PCR analysis confirmed the CCNI2 up-regulation in patient $CD43^{+}CD11b^{+}CD14^{-}$ cells sorted at day 16 of culture. Shown are the averages of 3 independent experiments as mean±SD, **P<0.01; unpaired t-test. D) $CD34^{+}CD43^{+}$ cells derived from patient iPSC (D14 of culture) were transduced with lentiviruses encoding shSCR and shCCNI2, sorted on $GFP^{+}$ 2 days after transduction (Day 16 of culture) and cultured for 7 days in the presence of G-CSF, SCF and IL-3. CCNI2 inhibition measured by qRT-PCR (*P<0.05, unpaired t-test) (i) led to inhibition of proliferation (ii) confirming that CCNI2 up-regulation in patient cells contributes to the increased proliferation rate. Shown are the averages of 3 independent experiments (n=1 for shCCNI2_1 and n=2 for shCCNI2_2) as mean±SD, ***P<0.005; ****P<0.001; 2-way ANOVA with multiple comparisons.

**Supplementary Figure 7**

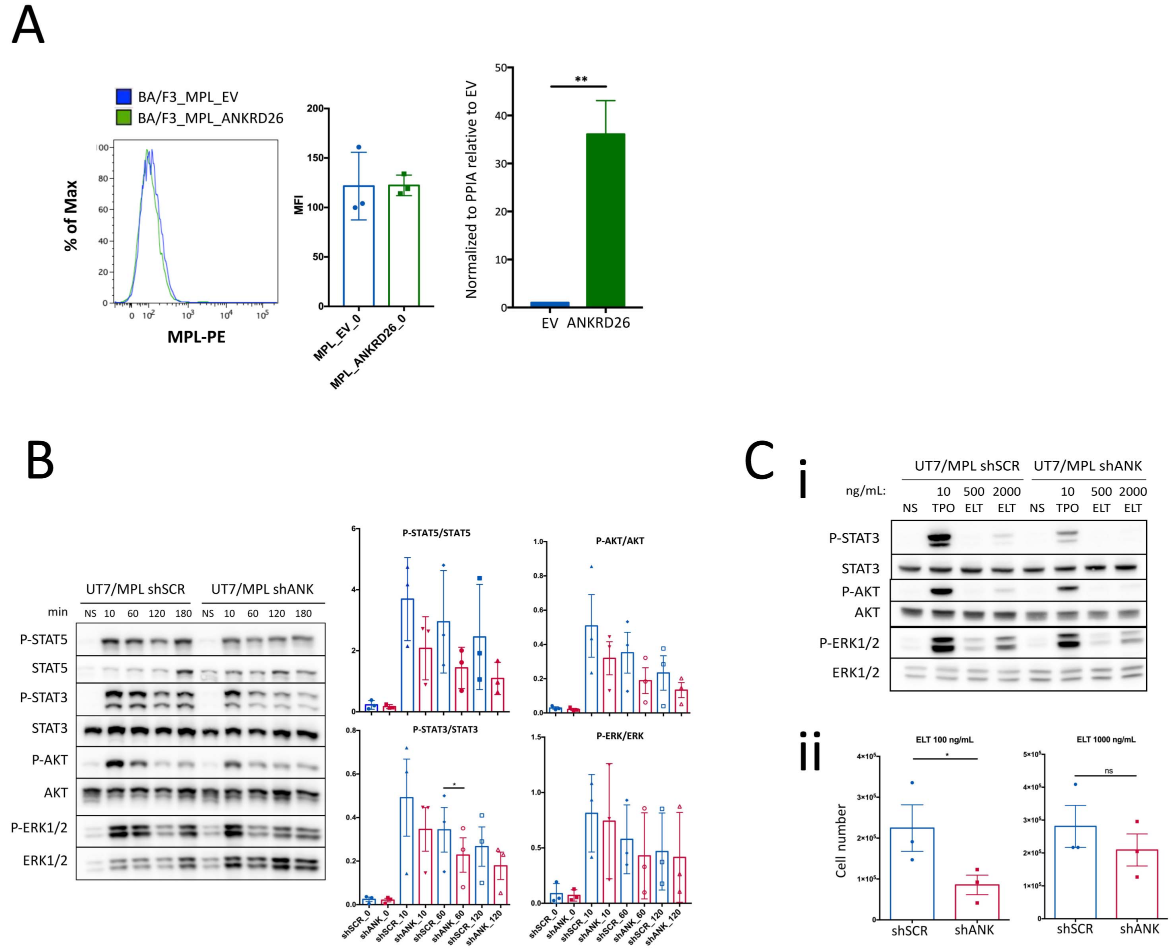


**Supplementary Figure 7. Higher ANKRD26 leads to the increased TPO/MPL and ELT/MPL signaling.**

A) HA_MPL expressing Ba/F3 (Ba/F3_MPL) cell line was transduced with a lentivirus harboring the cDNA of ANKRD26_V5 (ANKRD26) or with an empty lentivirus (EV) and transduced cells were selected by hygromycin B. No difference in MPL expression at the cell surface was detected after over-night starvation of cell as measured with an anti-MPL-PE antibody. Human *ANKRD26* transcript level was normalized to *PPIA* housekeeping gene and compared to the empty virus, condition where the arbitrary value being 1 as human ANKRD26 is not expressed in murine Ba/F3 cells.

B-C) HA_MPL expressing UT7 cell line (UT7/MPL) was transduced with the lentiviruses harboring control scramble shRNA (shSCR) or shANK (shANK_1 or shANK_2). C) Western blot analysis and quantification of TPO induced signaling in UT7/HA_MPL (UT7/MPL) cells transduced with shSCR or shANK at different times post-stimulation of cells with 10 ng/mL of TPO. The histograms show quantification of the WBs representing averages of 3 independent experiments as mean±SD. *P<0.05; paired t-test. C) Western blot analysis at different doses of Eltrombopag (ELT) at 10 min of stimulation showing that higher ANKRD26 expression (shSCR) leads to a hypersensitivity of MPL to ELT (i). Cell number of UT7/MPL cells measured at day 4 of culture with 2 different ELT doses is shown as mean±SD of 3 independent experiments (2 with shANK_1 and 1 with shANK_2), *P<0.05, ns: non-significant, paired t-test with was used (ii).

**Supplementary Figure 8**

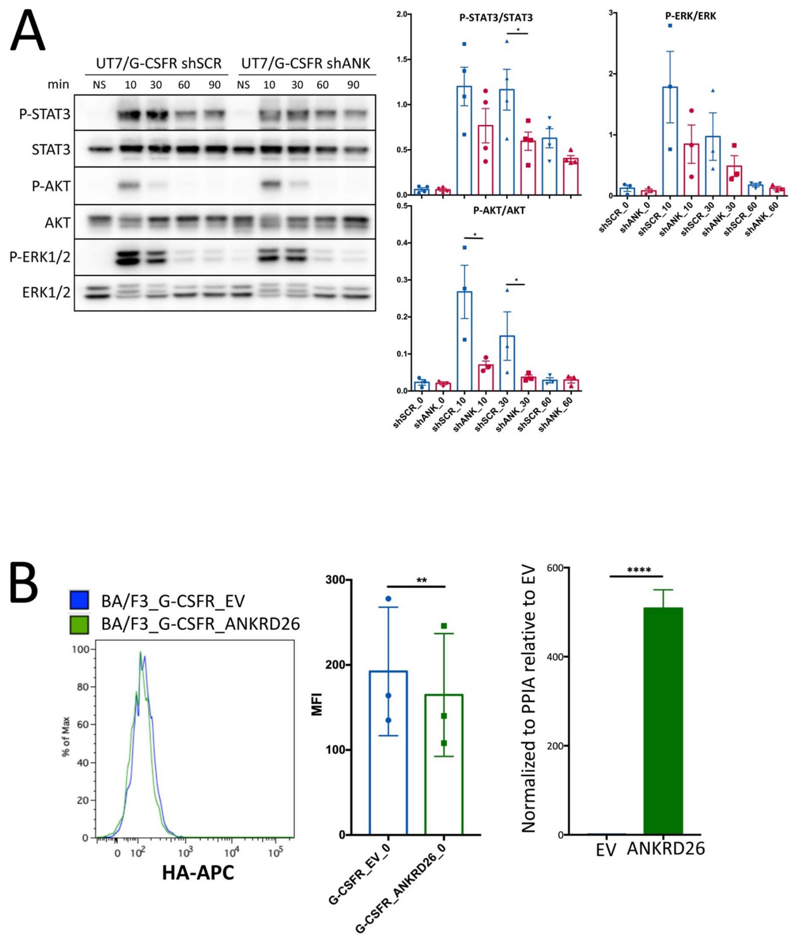


**Supplementary Figure 8. Higher ANKRD26 leads to the increased G-CSF/G-CSFR signaling.**

A) Western blot analysis and quantification of G-CSF induced signaling in UT7/HA_G-CSFR (UT7/G-CSFR) cells transduced with shSCR or shANK (shANK_1 or shANK_2) at different times post-stimulation of cells with 20 ng/mL of G-CSF. The histograms show quantification of the WBs representing averages of 3 or 4 independent experiments as mean±SD. *P<0.05; paired t-test. B) HA_G-CSFR expressing Ba/F3 (Ba/F3_G-CSFR) cell line was transduced with a lentivirus harboring the cDNA of ANKRD26_V5 (ANKRD26) or with an empty lentivirus (EV) and transduced cells were selected by hygromycin B. No difference in G-CSFR expression at the cell surface was detected after over-night starvation of cell as measured with an anti-HA antibody. Human *ANKRD26* transcript level was normalized to *PPIA* housekeeping gene and compared to the empty virus.

**Supplementary Figure 9**

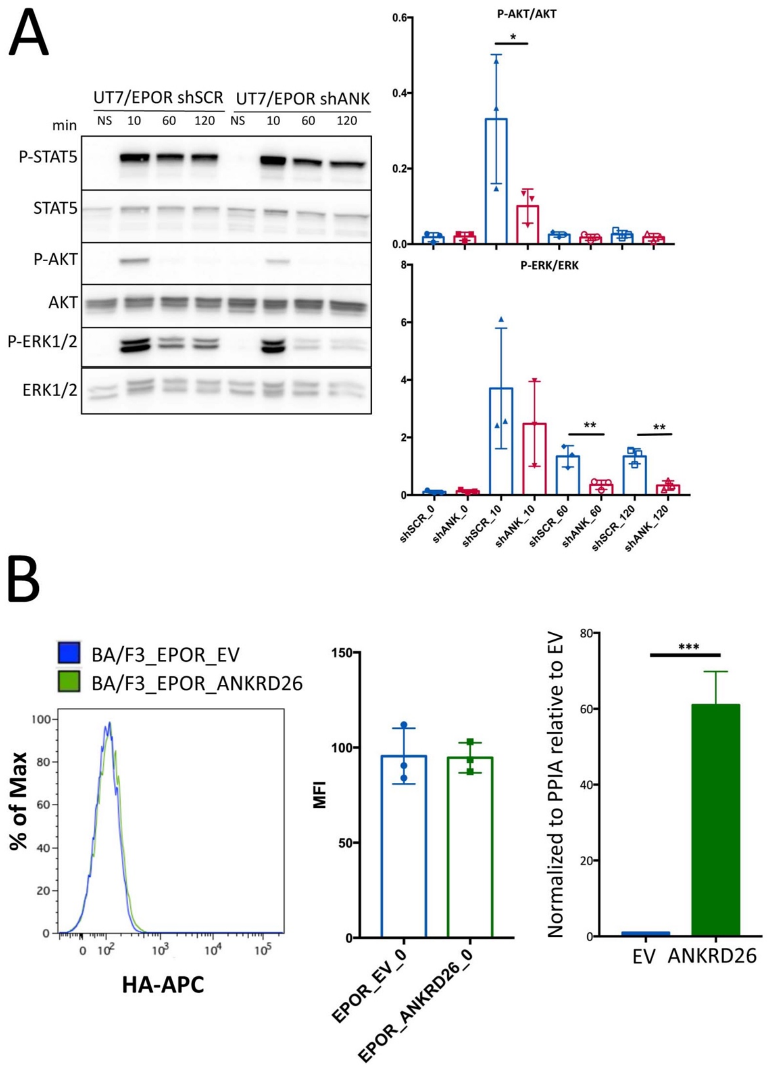


**Supplementary Figure 9. Higher ANKRD26 leads to the increased EPO/EPOR signaling.** A) Western blot analysis and quantification of EPO induced signaling in UT7/HA_EPOR (UT7/EPOR) cells transduced with shSCR or shANK (shANK_1 or shANK_2) at different times post-stimulation of cells with 1 U/mL of EPO. The histograms show quantification of the WBs representing averages of 3 independent experiments as mean±SD. *P<0.05; paired t-test. B) HA_EPOR expressing Ba/F3 (Ba/F3_EPOR) cell line was transduced with a lentivirus harboring the cDNA of ANKRD26_V5 (ANKRD26) or with an empty lentivirus (EV) and transduced cells were selected by hygromycin B. No difference in EPOR expression at the cell surface was detected after over-night starvation of cell as measured with an anti-HA antibody. Human *ANKRD26* transcript level was normalized to *PPIA* housekeeping gene and compared to the empty virus.

**Supplementary Figure 10**

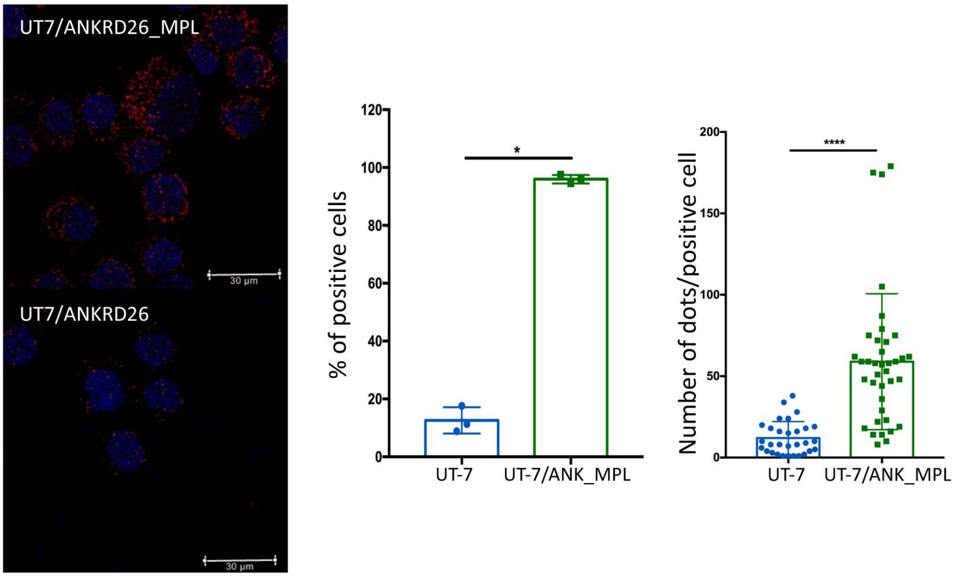


**Supplementary Figure 10. ANKRD26 and MPL interaction**

Representative pictures of proximity ligation assay for the ANKRD26 and MPL interaction in UT7 cell line. FLAG_ANKRD26 (ANK) and HA_MPL (MPL) were overexpressed in UT7 cell line. Monoclonal anti-FLAG antibody was used for ANKRD26 and polyclonal anti-HA for MPL. The representative pictures (red staining, left panel) represent the interaction between ANKRD26 and MPL, scale bar = 30 µm. Data (middle panel) represent the mean±SD of 3 independent experiments. At least 40 positive cells were analyzed for each condition (right panel). ***P<0.005, ****P<0.001, t-test with Mann Whitney correction.

### Supplementary Table 1. List of primers, antibodies and other products used

| Primer | Sequence | Experiment |
|---|---|---|
| ANKRD26_F | CTATGTCAGAGGCTTCACTGGAG | qPCR |
| ANKRD26_R | CTCAGCACATCTGACAGCTTCTG | qPCR |
| PPIA_F | GTCAACCCCACCGTGTTCTT | qPCR |
| PPIA_R | CTGCTGTCTTTGGGACCTTGT | qPCR |
| SOX2_F | GACAGAGCCCATTTTCTCCA | qPCR |
| SOX2_R | AAATCCTGTCCTCCCATTCC | qPCR |
| NANOG_F | TATGCCTGTGATTTGTGGGC | qPCR |
| NANOG_R | GTTTGCCTTTGGGACTGGTG | qPCR |
| OCT-4_F | AGAGGATCACCCTGGGATAT | qPCR |
| OCT-4_R | CGCCGGTTACAGAACCACAC | qPCR |
| GAPDH_F | TGTTCGACAGTCAGCCGCAT | qPCR |
| GAPDH_R | TAAAAGCAGCCCTGGTGACC | qPCR |
| CCNI2_F | TGACTATATTCCATGCCCTGGTG | qPCR |
| CCNI2_R | AGTGTGGAGCCCTTGAACTG | qPCR |
| seq_ANKRD26_R1 | AGCCATTCTTCCTGAGCAAA | sequencing |
| seq_ANKRD26_R2 | TTGTGGGAAGGTTTTGGAAG | sequencing |
| seq_ANKRD26_F3 | CTTCCAAAACCTTCCCACAA | sequencing |
| seq_ANKRD26_R4 | AGTTGCCTCTGAGCGTCATT | sequencing |
| seq_ANKRD26_R5 | TGCAGTCTCCAGTTCTGCTG | sequencing |
| seq_ANKRD26_F6 | TTACAGGCACAAGCAGCATC | sequencing |
| seq_ANKRD26_R7 | GCATCATCCAGTTGTTGTCG | sequencing |
| ANKRD26_pEF6/V5-His_F | TACCGAGCTCGGATCCCACCATGAAGAAGATTTTTAGTAAG | InFusion cloning |
| ANKRD26_pEF6/V5-His_R | GCCACTGTGCTGGATGATCATATAATTTTTCTTTAAAACC | InFusion cloning |
| PRRL_ANK_F | TCCTAGGCCTACGCGCCACCATGAAGAAGATTTTTAGT | InFusion cloning |
| PRRL_ANK_R | GCATGCCGGCACGCGTCAATGGTGATGGTGATGATGA | InFusion cloning |
| Infusion_FLAG_ANKRD26_3e_F | TCCTAGGCCTACGCGTCCACCATGGACTACAAGGACGACGATGACAAGAAGAAGATTTTTAGTAAGAAG | InFusion cloning |
| Infusion_FLAG_ANKRD26_2e_R | GCATGCCGGCACGCGTTCAGATCATATAATTTTTCTTTAAAACCTGTAC | InFusion cloning |
| shCCNI2_1 | TACCTGCATTGCGCCACAATT | Lentiviral cloning |
| shCCNI2_2 | CTGGACTTCTTGACTATATTC | Lentiviral cloning |

| **Antibody** | **Source** | **Clone** |
|---|---|---|
| CD34-PE | BD Biosciences | 581 |
| CD34-PE-Cy7 | Biolegend | 581 |
| CD43-APC | BD Biosciences | 1G10 |
| CD11b-PE | BD Biosciences | ICRF44 |
| CD14-APC-H7 | BD Biosciences | M5E2 |
| CD15-V450 | BD Biosciences | HI98 |
| CD45-FITC | BD Biosciences | HI30 |
| Oct3/4 | Santa Cruz | C-10 |
| Sox2 | Millipore | Polyclonal |
| TRA 1-81 | Millipore | TRA-1-81 |
| SSEA4 | Millipore | MC-813-70 |
| Nanog | Abcam | Polyclonal |
| IgM-APC | BD Biosciences | G155-228 |
| IgG1-PE | BD Biosciences | MOPC-21 |
| IgG1-PerCP-Cy™5.5 | BD Biosciences | MOPC-21 |
| IgG2a-Alexa Fluor® 647 | BD Biosciences | MOPC-173 |
| IgM-Alexa Fluor® 488 | Invitrogen | Polyclonal |
| IgG-Alexa Fluor® 488 F(ab')2 fragment | Invitrogen | Polyclonal |
| IgG-Alexa Fluor® 568 F(ab')2 fragment | Invitrogen | Polyclonal |
| CD110-PE | BD Biosciences | 1.6.1 |
| CD110-APC | BD Biosciences | 1.6.1 |
| CD235/GPA-PE-Cy7 | BD Biosciences | GA-R2 (HIR2) |
| CD36-APC | BD Biosciences | CB38 (NL07) |
| HA-APC | Miltenyi | GG8-1F3.3.1 |
| STAT3 | Cell Signaling Technology | D1A5 |
| STAT5 | Cell Signaling Technology | D3N2B |
| AKT (pan) | Cell Signaling Technology | C67E7 |
| p42/44 MAPK (Erk1/2) | Cell Signaling Technology | Polyclonal |
| Phospho STAT3 (Y705) | Cell Signaling Technology | D3A7 |
| Phospho STAT5 (Y694) | Cell Signaling Technology | D47E7 |
| Phospho AKT (S473) | Cell Signaling Technology | D9E |
| Phospho p42/44 MAPK (Erk1/2) (T202/Y204) | Cell Signaling Technology | Polyclonal |
| Actin | Sigma | AC-15 |
| IgG anti-rabbit H+L Alexa Fluor 633 | Invitrogen/Thermo Fisher Scientific | Polyclonal |
| IgG anti-mouse H+L Alexa Fluor 546 | Invitrogen/Thermo Fisher Scientific | Polyclonal |
| IgG anti-rabbit HRP-linked | Cell Signaling Technology | Polyclonal |
| IgG anti-mouse HRP-linked | Cell Signaling Technology | Polyclonal |
| TPOR/c-MPL | Millipore | Polyclonal |
| FLAG rabbit | Millipore, Sigma | Polyclonal |
| V5 mouse | Invitrogen | Unknown clone |
| HA mouse | Biolegend | 16B12 |
| CD4-APC | BD Pharmingen | 555349 |

| | | |
|---|---|---|
| Annexin -PE | BD Pharmingen | 5556421 |
| HA rabbit | Abcam | Polyclonal |
| CD41a-APC | BD Biosciences | HIP8 |
| CD42-PE | BD Biosciences | ALMA.16 |
| **Reagent** | **Source** | **Identifier** |
| hBMP4 | Peprotech | AF-120-05ET |
| hVEGF | Peprotech | 100-20 |
| hFGF-basic | Peprotech | 100-18B |
| hIL-6 | Peprotech | 200-06 |
| hEPO | Peprotech | 100-64 |
| hG-CSF | Peprotech | 300-23 |
| hGM-CSF | Peprotech | 300-03 |
| hIL-3 | Peprotech | 200-03 |
| hTPO | Kyowa Kirin, Tokyo, Japan | / |
| hFLT3l | Celldex Therapeutics, Inc., Needham, USA | / |
| hSCF | Biovitrum AB, Stockolm, Sweden | / |
| CHiR 99021 trihydrochloride | TOCRIS | 4953 |
| Y-27632 dihydrochloride | TOCRIS | 1254 |
| VTN-N | Gibco/Thermo Fisher Scientific | A14700 |
| Geltrex | Gibco/Thermo Fisher Scientific | A1413202 |
| StemPro®-34 SFM | Gibco/Thermo Fisher Scientific | 10639011 |
| E8 | Gibco/Thermo Fisher Scientific | A1517001 |
| E8 Flex | Gibco/Thermo Fisher Scientific | A2858501 |
| CytoTune™-iPS 2.0 Sendai Reprogramming Kit | Invitrogen/Thermo Fisher Scientific | A16517 |
| Minimum Essential Medium a | Gibco/Thermo Fisher Scientific | 22561021 |
| Fetal Bovine Serum | Hyclone | SV30160.03 |
| 1-thioglycerol | Sigma | M6145 |
| cOmplete™ inhibitor cocktail | Sigma/Roche | 4693159001 |
| Phosphatase Inhibitor Cocktail 3 | Sigma | P0044 |
| PMSF | Sigma | P7626 |
| Sodium Orthovanadate | Sigma | S6508 |
| Sodium Fluoride | Sigma | 201154 |
| Duolink® In Situ Red Starter Kit Mouse/Rabbit | Sigma | DUO92101 |
| DMEM | Gibco/Thermo Fisher Scientfic | 41966 |
| Trypsine/EDTA 0.05% | Sigma | 23500054 |
| TrypIE Express | Gibco/Thermo Fisher Scientfic | 12605036 |
| Ascorbic Acid | Gibco/Thermo Fisher Scientfic | A4403 |
| TransIT®-293 Transfection Reagent | Mirus Bio LLC | MIR 2706 |
| Penicillin/Streptomycin | Gibco/Thermo Fisher Scientfic | 15140-122 |
| Glutamin | Gibco/Thermo Fisher Scientfic | 25030123 |

**Supplementary Table 2**

| | Mutation | Sex | Family history of malignancy | Platelet count ($x10^9$/L) | Experiment (Figure) |
|---|---|---|---|---|---|
| **P1** | c. -127delAT | M | No | 55 | Figure 4Ai, ii, B, C, D, E |
| **P2** | c. -127delAT | M | No | 56 | Figure 4B, D |
| **P3** | c.-127A>T | M | No | 36 | Figures 4B, C; 7Dii |
| **P4** | c. -118C>A | M | No | 45 | Figure 4Ai, ii, B, C |
| **P5** | c.-126T>C | M | No | 20 | Figure 4Ai, 7Dii |
| **P6** | c.-116C>T | M | No | 69 | Figure 4Ai, 7Dii |

**Suppelementary Table 2:** List of patient samples used for experiments performed on primary cells

**Supplementary Table 3**

| | Mutation | Sex | Family history of malignancy | Platelet count ($x10^9$/L) | iPSC cell line |
|---|---|---|---|---|---|
| **P4** | c. -118C>A | M | No | 45 | ANK1 |
| **P7** | c. -127delAT | M | No | 26 | ANK2 |

**Supplementary Table 3**: List of patient samples used for iPSC lines generation.

**Supplementary Table 4**

| Upregulated genes | | |
|---|---|---|
| ID | logFC | p-Value |
| AC079466.1 | 3.07 | 3.39e-08 |
| ADARB2 | 2.48 | 3.36e-07 |
| APOL1 | 2.80 | 1.27e-07 |
| CCNI2 | 3.54 | 2.23e-12 |
| CD2 | 3.07 | 3.64e-11 |
| CD3D | 1.85 | 3.84e-06 |
| CES1P1 | 2.79 | 1.69e-07 |
| EPS8L3 | 2.29 | 5.62e-07 |
| GAPDHP1 | 2.61 | 5.34e-10 |
| GIMAP7 | 2.44 | 2.30e-07 |
| GPT | 2.47 | 1.30e-06 |
| HLA-DQA1 | 3.44 | 9.00e-12 |
| HLA-DQB1 | 3.66 | 6.78e-13 |
| IL13 | 2.48 | 5.76e-07 |
| NPTX1 | 2.61 | 1.66e-07 |
| POF1B | 2.42 | 1.10e-05 |
| RP11-188C12.2 | 3.18 | 1.64e-09 |
| RP11-455F5.3 | 2.41 | 2.92e-06 |
| RP4-737E23.2 | 2.33 | 5.43e-06 |
| SIGLEC12 | 2.93 | 3.29e-17 |
| SP7 | 2.69 | 2.89e-07 |
| SSXP10 | 2.39 | 1.39e-05 |
| SULT1B1 | 3.04 | 1.33e-09 |
| TNFRSF18 | 2.16 | 6.21e-07 |

| Downreguated genes | | |
|---|---|---|
| ID | logFC | p-Value |
| CCDC144NL-AS1 | -2.34 | 1.74e-07 |
| OSMR | -1.87 | 1.19e-05 |
| ZFP28 | -3.48 | 1.76e-14 |
| ZFP42 | -3.43 | 2.33e-10 |
| ZNF486 | -4.35 | 1.49e-16 |
| ZNF626 | -4.01 | 3.00e-14 |
| ZNF682 | -3.08 | 1.22e-08 |

**Supplementary Table 4.** List of up- and down-regulated genes in $CD43^{+}CD11b^{+}CD14^{-}$ progenitors derived from ANK1&2 iPSC lines versus control iPSC lines.
